# Peripheral metabolic dysfunction drives sleep disruption in TDP-43 proteinopathy

**DOI:** 10.64898/2026.02.23.707321

**Authors:** Anyara Rodriguez, Jenny Luong, Samuel J. Belfer, Oksana Shcherbakova, Jaclyn Durkin, Pinky Kain, Arjun Sengupta, Alexandra E. Perlegos, Farheen Akhtar, Daniel J. Boehmler, Zhecheng Jin, Aalim M Weljie, Nancy M. Bonini, Matthew S. Kayser

**Affiliations:** Department of Psychiatry, Perelman School of Medicine at the University of Pennsylvania, Philadelphia, PA, 19104, USA; Department of Biology, University of Pennsylvania, Philadelphia, PA, 19104, USA; Department of Systems Pharmacology and Translational Therapeutics, University of Pennsylvania, Philadelphia, PA, 19104, USA; Institute for Translational Medicine and Therapeutics, University of Pennsylvania, Philadelphia, PA, 19104, USA; Department of Cancer Biology, Perelman School of Medicine at the University of Pennsylvania, Philadelphia, PA, 19104, USA; Department of Radiology, Perelman School of Medicine at the University of Pennsylvania, Philadelphia, PA, 19104, USA; Chronobiology Sleep Institute, Perelman School of Medicine at the University of Pennsylvania, Philadelphia, PA, 19104, USA; Department of Neuroscience, Perelman School of Medicine at the University of Pennsylvania, Philadelphia, PA, 19104, USA

## Abstract

Sleep disruption is an early and prevalent feature of neurodegenerative disease, commonly attributed to neuronal circuit dysfunction or cell loss. However, sleep is tightly coupled to metabolic state, raising the possibility that systemic metabolic abnormalities contribute to disease-associated sleep phenotypes. Using *Drosophila* models of TDP-43 proteinopathy, we investigated whether peripheral metabolic dysfunction plays a causal role in sleep disruption. We show that TDP-43 expression induces a chronic, starvation-like metabolic state characterized by depletion of peripheral carbohydrate stores despite normal feeding. Restoration of sleep fails to correct these metabolic defects, whereas improving peripheral metabolic state robustly rescues sleep. A modifier screen of ∼650 RNAi lines identified Salt-inducible kinase 3 (SIK3) as a potent suppressor of both sleep loss and starvation sensitivity. Transcriptomic and spatial metabolomic analyses reveal that SIK3 selectively remodels a peripheral metabolic program centered on the pentose phosphate pathway and redox-associated metabolites without globally restoring energy stores. Together, these findings identify systemic metabolic dysfunction as a key driver of sleep disruption in TDP-43 proteinopathy and highlight peripheral metabolism as a potential therapeutic entry point for sleep dysfunction in neurodegenerative disease.

## Introduction

Sleep disruption is a prominent and often early feature of neurodegenerative disease, including amyotrophic lateral sclerosis (ALS) and frontotemporal dementia (FTD)^1–4^. Alterations in sleep duration and continuity are frequently observed prior to advanced cognitive or motor decline, suggesting that sleep dysfunction may reflect early disease biology rather than be a consequence of later neuronal loss^1^. At the same time, sleep plays a critical role in maintaining neuronal homeostasis, metabolic balance, and stress resilience^5–8^, raising the possibility that sleep disruption may not only mark disease onset but also contribute to disease progression.

In parallel with sleep disturbances, metabolic dysfunction is increasingly recognized as a core feature of ALS and FTD^9–13^. Patients exhibit alterations in energy balance, including hypermetabolism, weight loss, altered glucose utilization, and dyslipidemia; metabolic state can predict disease trajectory and survival. Importantly, these metabolic abnormalities are not restricted to the central nervous system: peripheral tissues show altered carbohydrate and lipid metabolism, and systemic metabolic stress can precede or exacerbate neurological decline^11,12^. Despite this, most mechanistic studies of ALS/FTD have focused on neuronal vulnerability and circuit dysfunction, leaving unresolved whether systemic metabolic abnormalities actively influence behavioral phenotypes such as sleep. This gap is notable given that TDP-43, while prominently studied in neurons, is expressed across cell types and tissues and that pathological TDP-43 abnormalities have been documented in peripheral tissues in disease^14,15^. Consistent with this possibility, systemic metabolic state is tightly coupled to sleep regulation across species^16–21^.

TDP-43 proteinopathy is a defining pathological feature of ALS, a large fraction of FTD, and other neurodegenerative diseases including Alzheimer’s^14,22–25^. TDP-43 is a multifunctional RNA-binding protein involved in transcription, splicing, RNA transport, and stress responses; in disease, TDP-43 becomes mislocalized, aggregated, and post-translationally modified, with profound consequences for cellular function^14,24^. Expression of human TDP-43 in *Drosophila* is toxic and recapitulates key ALS/FTD disease features, including neuronal degeneration and shortened lifespan^26–29^. In prior work, we showed that adult-onset expression of human TDP-43 in flies is associated with severe sleep loss and fragmentation, as well as brain metabolic dysregulation^30^. Ataxin-2, a genetic suppressor of TDP-43 neurodegeneration, restored sleep and normalized brain glycogen levels^30^. These findings link the TDP-43-associated sleep disruption to an altered metabolic state in the nervous system. However, whether TDP-43–induced sleep disturbances arise from brain-intrinsic metabolic dysfunction or whether systemic metabolic abnormalities play a causal role is not known.

Here, we find that human TDP-43 expression in *Drosophila* induces a systemic, starvation-like metabolic state characterized by depletion of peripheral carbohydrate stores, despite normal feeding behavior. We further demonstrate that restoring sleep fails to correct these metabolic defects, whereas improving peripheral metabolic state through dietary or genetic means can robustly rescue sleep. Through an RNAi-based enhancer/suppressor screen of the sleep phenotype, we identify Salt-inducible kinase 3 (SIK3) as a novel modifier of TDP-43. Behavioral analyses, transcriptomic profiling, and spatial metabolomics demonstrate that Sik3 knockdown suppresses both sleep loss and starvation sensitivity in TDP-43 flies without globally restoring energy stores. Instead, SIK3 selectively remodels a peripheral metabolic program centered on the pentose phosphate pathway and redox-associated metabolites. Together, these findings reveal that peripheral metabolic dysfunction is a key driver of sleep disruption in TDP-43 proteinopathy, extending models of sleep regulation in neurodegenerative disease beyond brain-intrinsic mechanisms.

## Results

### TDP-43 expression induces peripheral metabolic dysfunction

Sleep disruption is often accompanied by metabolic abnormalities, and metabolic stress can alter sleep behavior^31^. Our prior work focused on metabolic changes within the brain^30^, but TDP-43–expressing flies are also unusually sensitive to starvation^32^, indicating that systemic energy homeostasis may be compromised. We therefore determined whether TDP-43 proteinopathy induces peripheral metabolic defects. To assess global metabolic resilience, we examined survival under nutrient deprivation. Using the hormone-inducible GeneSwitch driver under control of the ubiquitous *daughterless* promoter (*Da*GS) to achieve broad, adult-restricted expression of UAS-*TDP-43^52S^*, we confirmed that TDP-43 flies exhibited markedly reduced starvation resistance compared to controls (Fig. 1A). This effect raised the possibility of impaired energy storage or utilization. We next directly quantified key metabolites in the bodies of animals expressing TDP-43. In contrast to unchanged glucose and an increase in glycogen found in brains of these animals^30^, whole-body assays revealed dramatic reductions in glucose and glycogen in TDP-43 flies (Fig. 1B), indicating depletion of both carbohydrate stores. To determine whether reduced energy stores reflected altered feeding behavior, we measured food intake using the CAFÉ assay^33^ and found that TDP-43 flies did not show a detectable reduction in food consumption compared to controls (Fig. S1); this argues against altered feeding as the primary driver of energy depletion. Instead, these data suggest that TDP-43-expressing flies are defective in metabolic storage, utilization, or routing.

**Figure 1.**
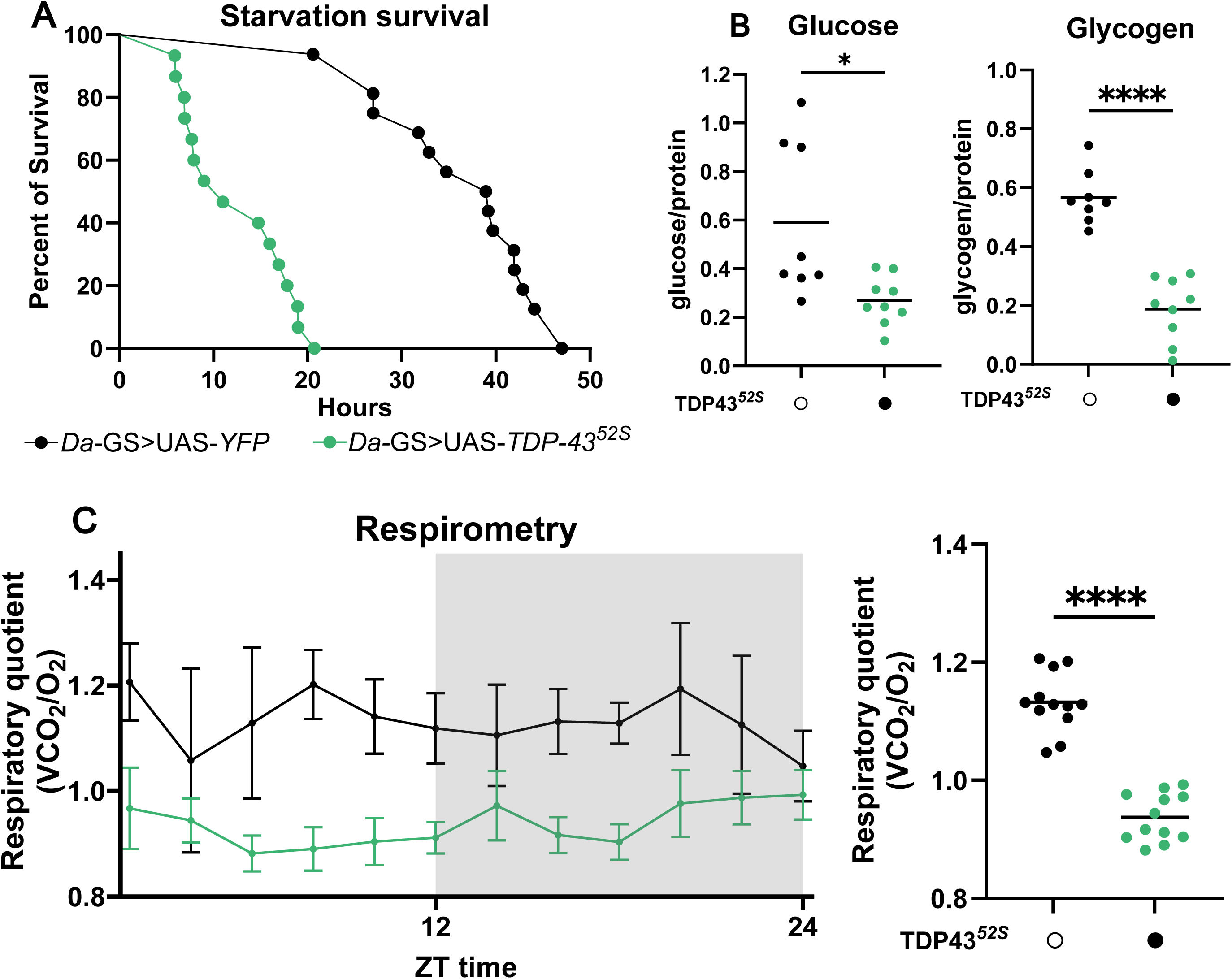
TDP-43 expression induces peripheral metabolic dysfunction. (**A**) Reduced survival of TDP-43 flies (n=15 flies) compared to control flies (n=16 flies) under starvation. \*\*\*\**p* ≤ 0.0001 log-rank test. (**B**) Reduced glucose, glycogen levels in TDP-43 bodies (n=9, 9, 10 respectively) compared to control bodies (n=8, 8, 12 respectively). Each data point represents 5 bodies, \**p* ≤ 0.05, \*\**p* ≤ 0.01, and \*\*\*\**p* ≤ 0.0001, unpaired t-test. (**C**) Respiratory quotient (RQ = VCO₂/VO₂) in control and TDP-43 flies measured over a 24 h recording period. RQ values were binned into 2 h intervals and plotted as mean ± SEM across one complete light:dark cycle (left; gray shading indicates the dark phase) or averaged over the 24 h period (right). \*\*\*\**p* ≤ 0.0001, one-way ANOVA with Dunnett’s multiple comparisons test.

To assess whole-animal substrate utilization, we performed indirect calorimetry to measure carbon dioxide production (VCO₂) and oxygen consumption (VO₂) over a 24 h light:dark cycle and calculated the respiratory quotient (RQ = VCO₂/VO₂)^34,35^. RQ provides an index of fuel preference, with values closer to 1 indicating carbohydrate oxidation and lower values reflecting increased reliance on lipid or protein substrates^35^. TDP-43 flies exhibited a consistently lower RQ than controls across the recording period, indicating reduced reliance on carbohydrate metabolism and increased use of alternative energy sources (Fig. 1C). This shift is consistent with the observed depletion of peripheral carbohydrate stores and supports the idea that TDP-43 flies exist in a chronic starvation-like metabolic state despite continued access to food.

We next compared transcriptomic profiles of TDP-43 expression in brains and peripheral tissues (bodies) to assess whether molecular consequences differed between these regions (Fig. 2A). TDP-43 expression produced widespread transcriptional remodeling in both tissues, with ∼6,000 transcripts differentially expressed in bodies and >3,000 in brains relative to controls, including a shared subset of 1821 transcripts altered in both contexts (Fig. 2B). Thus, TDP-43 elicits both common and tissue-specific transcriptional responses. To determine whether particular biological programs were differentially affected, we performed pathway-level enrichment analyses stratified by tissue and direction of transcriptional change (Fig. 2C, Fig. S2A-C). As previously reported in brain^30^ and now observed in bodies (Fig. S2), TDP-43 expression resulted in upregulation of many common metabolic pathways/processes. However, differences emerged in the down-regulated gene sets. In the periphery, transcripts with reduced expression were selectively enriched for core metabolic programs that included glucose metabolism, glycolysis/gluconeogenesis, carbon metabolism, and the pentose phosphate pathway (PPP); these enrichments were absent in brains (Fig. 2C, Fig. S2). This pattern supports a suppression of peripheral metabolic capacity in TDP-43 animals that is distinct from transcriptional metabolic changes in the brain. Together, these data demonstrate depleted peripheral carbohydrate energy stores in TDP-43 flies, consistent with a chronic starvation-like physiology despite normal feeding.

**Figure 2.**
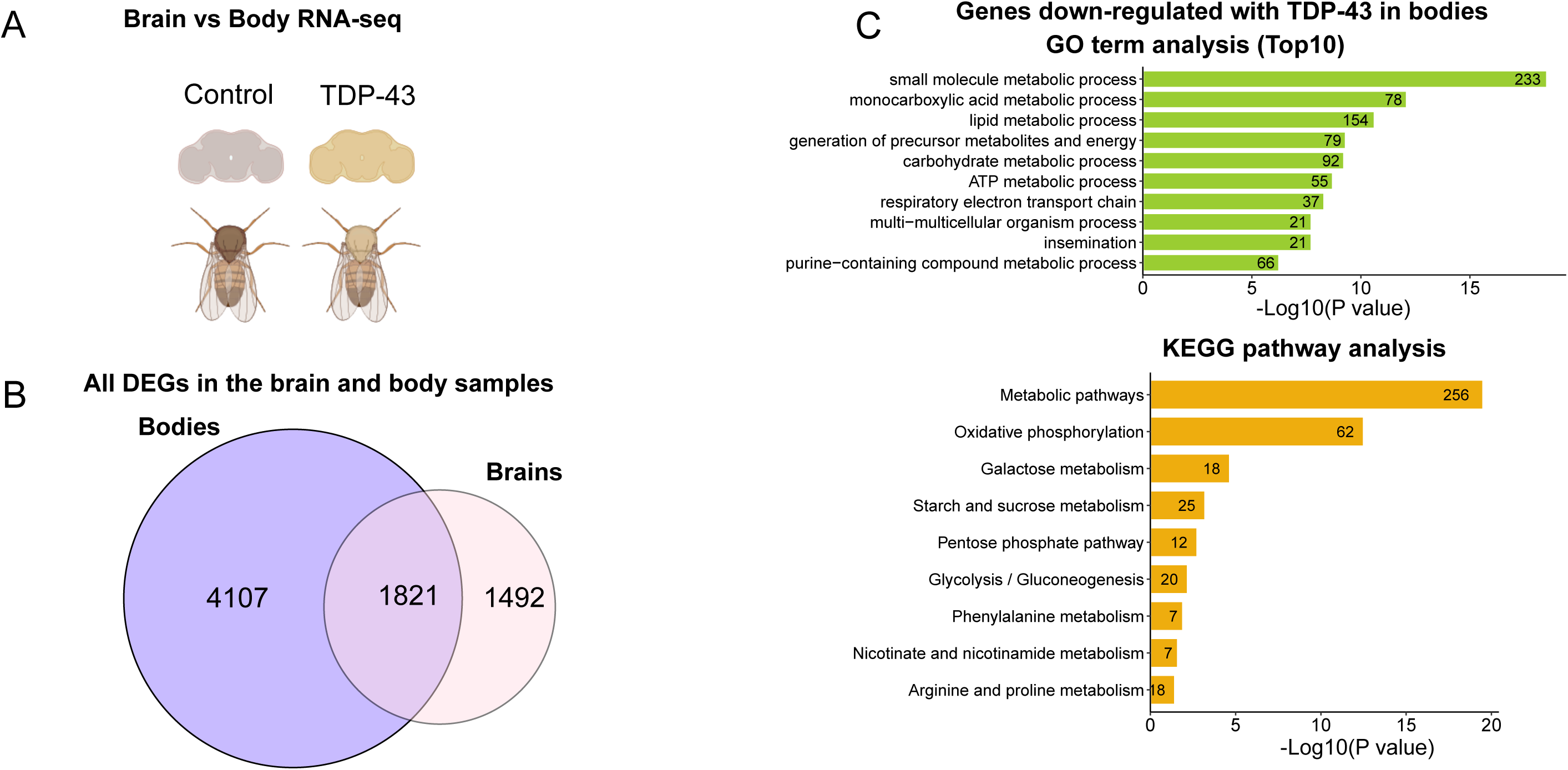
Comparison of RNA-seq from brains and bodies of TDP-43 animals. (**A**) Experimental design. Brain dataset from Perlegos et al, 2024^30^. Bodies of genotype daGS; UAS-YFP x Luc-RNAi or daGS; UAS-TDP-43 x Luc-RNAi were processed for RNA-seq following 6d on RU486. (**B**) Differentially expressed transcripts in bodies and brains of TDP-43 flies. (**C**) Gene ontology (GO) term and Kyoto Encyclopedia of Genes and Genomes (KEGG) pathway analysis of genes down-regulated in TDP-43 bodies.

### Inducing sleep does not normalize peripheral metabolic defects in TDP-43 flies

Sleep loss itself can disrupt metabolic homeostasis^31^, raising the possibility that peripheral metabolic abnormalities in TDP-43 flies are downstream of the sleep disruption. To address this directionality, we asked whether pharmacologically restoring sleep in TDP-43 flies was sufficient to normalize peripheral metabolism. We induced sleep in adult flies using gaboxadol, a GABAA receptor agonist that robustly increases sleep^36^. Gaboxadol treatment significantly increased total sleep time in TDP-43 flies and improved sleep consolidation (Fig. 3A; Fig S3).

**Figure 3.**
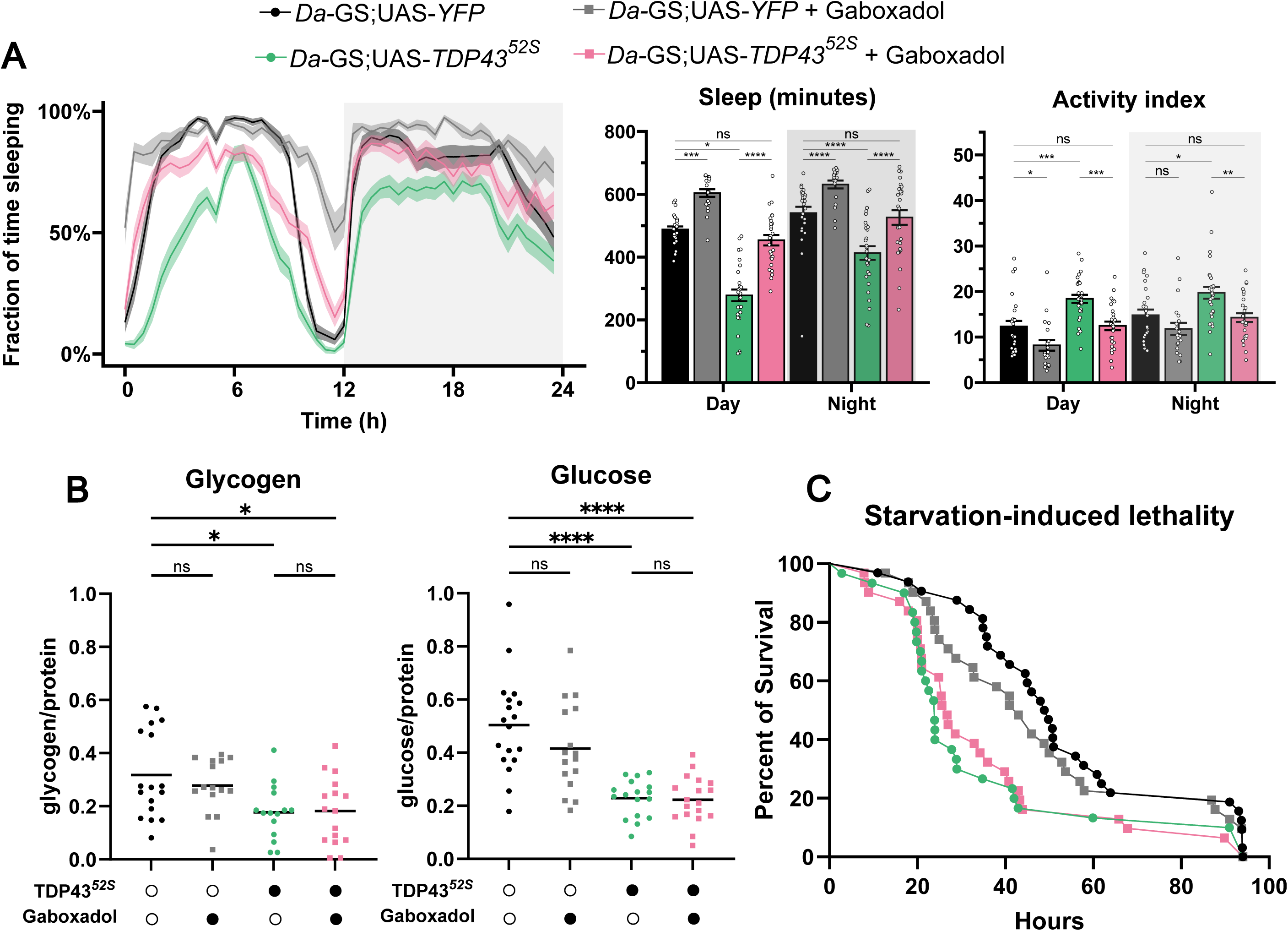
Inducing sleep with gaboxadol fails to normalize peripheral metabolic defects in TDP-43 flies. (**A**) Sleep traces over 24 hours and quantification of sleep duration and activity during wakefulness during the day and night. From left to right: control flies (n=29 flies), control flies + gaboxadol (n=20 flies), *TDP-43* flies (n=25 flies), *TDP-43* flies + gaboxadol (n=27 flies). \**p* ≤ 0.05, \*\**p* ≤ 0.01, *** *p* ≤ 0.001, and \*\*\*\**p* ≤ 0.0001 by one-way ANOVA with Šídák’s multiple comparisons test. (**B**) Glucose and glycogen levels in experimental and control bodies, each point represents 5 bodies. From left to right, for glucose assay n=18, 17, 16, 18, for glycogen assay n=18, 17, 16, 18. \**p* ≤ 0.05, \*\**p* ≤ 0.01, *** *p* ≤ 0.001, and \*\*\*\**p* ≤ 0.0001 by one-way ANOVA with Šídák’s multiple comparisons test. (**C**) Survival under starvation of *TDP-43* flies (23.9 hours, n=30 flies), control flies (49.3 hours, n=32 flies), *TDP-43* flies + gaboxadol (26.7 hours, n=31 flies), control flies + gaboxadol (43 hours, n=31 flies). *TDP-43* versus control: ns, *TDP-43* versus *TDP-43* + gaboxadol: ns, log-rank test with Holm-Šídák’s multiple comparisons test.

Despite effective sleep induction, peripheral metabolic defects persisted. Whole-body glucose and glycogen levels remained significantly reduced in TDP-43 flies treated with gaboxadol (Fig. 3B). Moreover, gaboxadol-treated TDP-43 flies remained highly sensitive to starvation (Fig. 3C), even though the drug was withdrawn prior to starvation assays, to exclude acute pharmacological effects. These results indicate that sleep disruption alone does not account for the peripheral metabolic phenotype in TDP-43 flies. Instead, metabolic dysfunction appears to occur upstream of, or in parallel with, sleep loss.

### Dietary glucose rescues sleep and peripheral metabolic defects in TDP-43 flies

We next asked whether improving metabolic state in TDP-43 flies could restore sleep. To address this, we switched adult flies to a high-glucose diet and examined sleep and metabolic outcomes. Dietary glucose supplementation significantly improved sleep in TDP-43 flies, increasing total sleep time and restoring sleep architecture (Fig. 4A; Fig. S4). Although control flies also exhibited trends towards increased sleep duration on high glucose, the magnitude of rescue in TDP-43 flies was substantially greater, consistent with a preferential benefit to these animals which are under metabolic stress. Moreover, high-glucose feeding restored peripheral glucose levels in TDP-43 flies (Fig. 4B) and rescued starvation sensitivity (Fig. 4C), demonstrating functional metabolic rescue. Thus, restoring peripheral glucose availability is sufficient to rescue sleep deficits in TDP-43 flies, linking metabolic state to sleep regulation.

**Figure 4.**
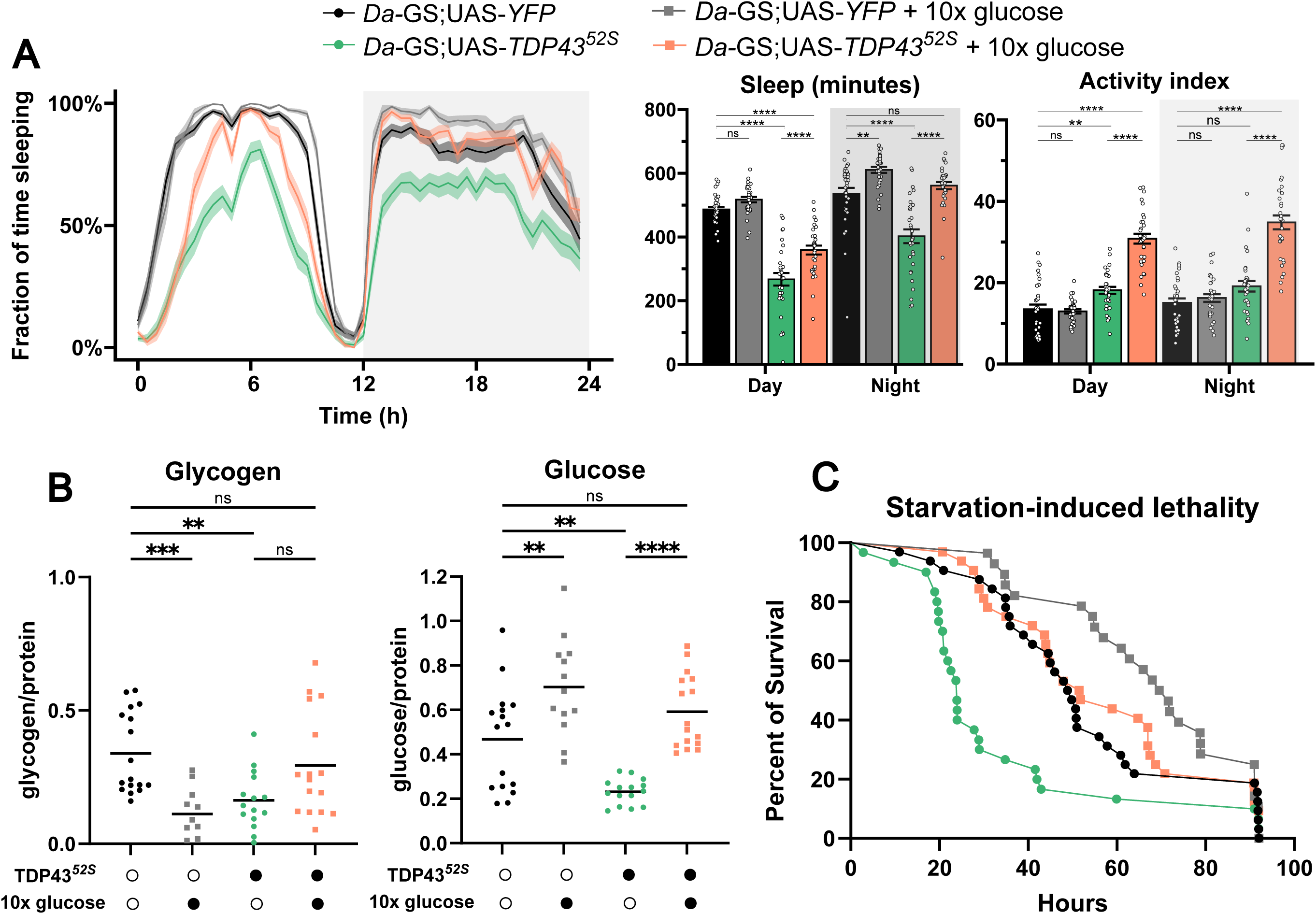
Dietary glucose rescues sleep and peripheral metabolic defects in TDP-43 flies. (**A**) Sleep traces over 24 hours and quantification of sleep duration and activity during wakefulness during the day and night. From left to right: control flies (n=32 flies), control flies + 10x glucose (n=31 flies), *TDP-43* flies (n=31 flies), *TDP-43* flies + 10x glucose (n=32 flies). \**p* ≤ 0.05, \*\**p* ≤ 0.01, *** *p* ≤ 0.001, and \*\*\*\**p* ≤ 0.0001 by one-way ANOVA with Šídák’s multiple comparisons test. (**B**) Glucose and glycogen levels in experimental and control bodies, each data point represents 5 bodies. From left to right, for glucose assay n= 19, 16, 18, for glycogen assay n= 20, 14, 18, 19 from left to right. \**p* ≤ 0.05, \*\**p* ≤ 0.01, *** *p* ≤ 0.001, and \*\*\*\**p* ≤ 0.0001 by one-way ANOVA with Šídák’s multiple comparisons test. (**C**) Survival under starvation of *TDP-43* flies (n=30 flies), control flies (n=32 flies), *TDP-43* flies + 10x glucose (n=32 flies), control flies + 10x glucose (n=28 flies). *TDP-43* versus control: *p* ≤ 0.05, *TDP-43* versus *TDP-43* + 10x glucose: *p* ≤ 0.01, log-rank test with Holm-Šídák’s multiple comparisons test.

### A genetic screen identifies SIK3 as a modifier of TDP-43 sleep and starvation phenotypes

To identify molecular pathways linking sleep with TDP-43 dysfunction, we performed an RNAi-based genetic screen of candidate modifiers of the sleep phenotype. Screening was conducted in *Da*GS > TDP-43 flies, covering ∼650 RNAi lines targeting genes either already implicated in TDP-43 pathology or sleep/circadian regulation and function (Fig. 5A; Table S1). Of the RNAi lines assayed in the primary screen, 141 resulted in enhancement or suppression of the TDP-43 sleep phenotype. Atx2, a well-characterized suppressor of TDP-43 toxicity^37^, was included as a positive control and strongly attenuated the sleep phenotype with knockdown, as previously shown^30^ (Fig. 5A). Interestingly, knockdown of several genes implicated in TDP-43 toxicity, such as *Chd1*^32^, failed to modify sleep, suggesting that the mechanisms of sleep pathology in TDP-43 flies may be distinct from other toxicity phenotypes. Among novel TDP-43 modifiers, knockdown of Salt-inducible kinase 3 (Sik3) emerged as a robust suppressor of TDP-43–induced sleep loss, with three independent RNAi lines targeting Sik3 exhibiting rescue of the TDP-43 sleep phenotype (Fig. 5A).

**Figure 5.**
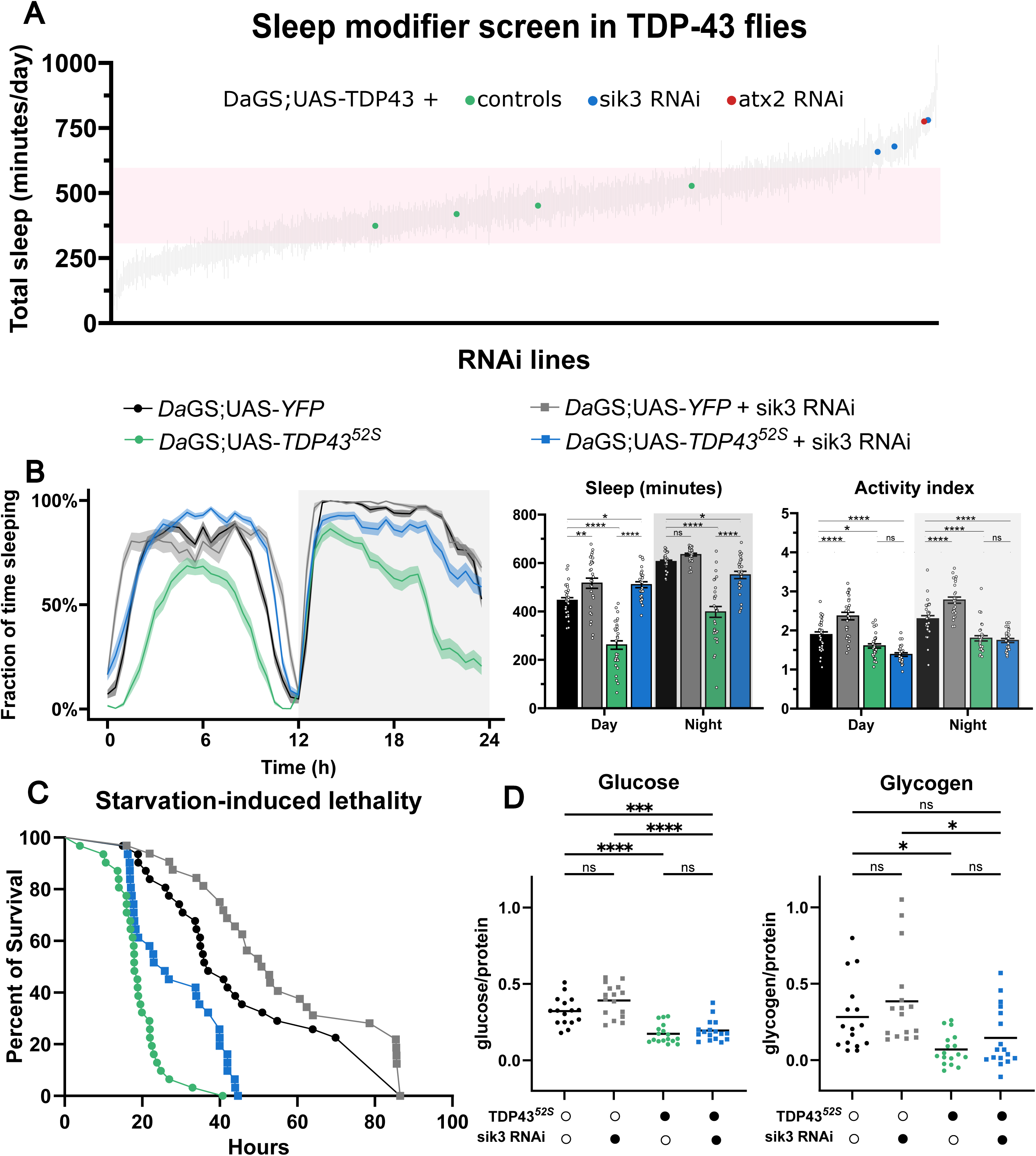
A genetic screen identifies SIK3 as a modifier of TDP-43 sleep and starvation phenotypes. (**A**) Sleep screen of RNAi lines co-expressed with *TDP-43* (green points are different RNAi controls with *TDP-43*). Pink box denotes one standard deviation from the mean sleep duration of all lines. (**B**) Sleep traces over 24 hours and quantification of sleep duration and activity during wakefulness during the day and night. From left to right: control flies (n=31), *TDP-43* flies (n=31), control flies + *sik3* RNAi (n=28), *TDP-43* flies + *sik3* RNAi (n=27). \**p* ≤ 0.05, \*\**p* ≤ 0.01, *** *p* ≤ 0.001, and \*\*\*\**p* ≤ 0.0001 by one-way ANOVA with Šídák’s multiple comparisons test. (**C**) Survival under starvation of *TDP-43* flies (n=31 flies), control flies (n=31 flies), *TDP-43* flies + *sik3* RNAi (n=31 flies), control flies + *sik3* RNAi (n=32 flies). \*\*\*\**p* ≤ 0.0001 log-rank test with Holm-Šídák’s multiple comparisons test. (**D**) Glucose and glycogen levels in *TDP-43* and control bodies with or without sik3 RNAi, each point represents 5 bodies. From left to right, for glucose assay n = 16, 16, 17, 17, for glycogen assay n = 16, 16, 17, 17. \**p* ≤ 0.05, \*\**p* ≤ 0.01, *** *p* ≤ 0.001, and \*\*\*\**p* ≤ 0.0001 by one-way ANOVA with Šídák’s multiple comparisons test.

Sik3, a member of the AMPK-related kinase family, is involved in metabolism and energy homeostasis^38,39^, as well as sleep regulation^19,40^. Reducing Sik3 expression restored total sleep time and improved sleep consolidation in TDP-43 flies, without producing strong sleep effects in control flies (Fig. 5B; Fig S5). qPCR validated efficient Sik3 knockdown (Fig. S6A) and we confirmed that Sik3 RNAi had no impact on TDP-43 expression levels (Fig S6B).

We then assessed general metabolic measures in TDP-43 flies with Sik3 knockdown and observed a reduction in starvation sensitivity (Fig. 5C), linking sleep rescue by Sik3 depletion to improved metabolic resilience. Surprisingly, despite this finding and normalized sleep in these flies, body glucose and glycogen levels were not consistently rescued (Fig. 5D). This apparent dissociation suggests that the suppression of sleep loss by Sik3 is not mediated by replenishing peripheral energy stores; instead, this result points toward the possibility of changes in metabolic routing and redox that could influence physiology and sleep, without restoring bulk glucose/glycogen content.

### TDP-43 sleep impairment is associated with peripheral PPP/redox dysfunction that is modified by Sik3 knockdown

As noted, TDP-43 proteinopathy produces a systemic starvation-like state characterized by reduced peripheral energy stores and impaired metabolic physiology, and dietary glucose can improve both sleep and starvation resistance. To identify metabolic processes most strongly altered by TDP-43 and corrected by Sik3 knockdown, we performed transcriptomic profiling of fly bodies and brains across genotypes (control, TDP-43, Sik3 RNAi, and TDP-43 + Sik3 RNAi; Fig. 6A). Principal component analysis highlighted tissue-specific differences. In bodies, TDP-43 expression produced a pronounced shift along PC1 relative to controls, reflecting widespread transcriptional remodeling (Fig. 6B). Notably, TDP-43 + Sik3 RNAi samples remained separated from controls along this axis, and thus were not reverted to a normal state, while Sik3 RNAi-only samples clustered near controls (Fig. 6B). In brains, separation was less pronounced and Sik3 knockdown did not substantially alter the brain transcriptome (Fig. 6C). These patterns indicate that Sik3 does not globally normalize TDP-43–induced transcriptional changes in bodies, but instead induces a distinct peripheral transcriptional state with TDP-43.

**Figure 6.**
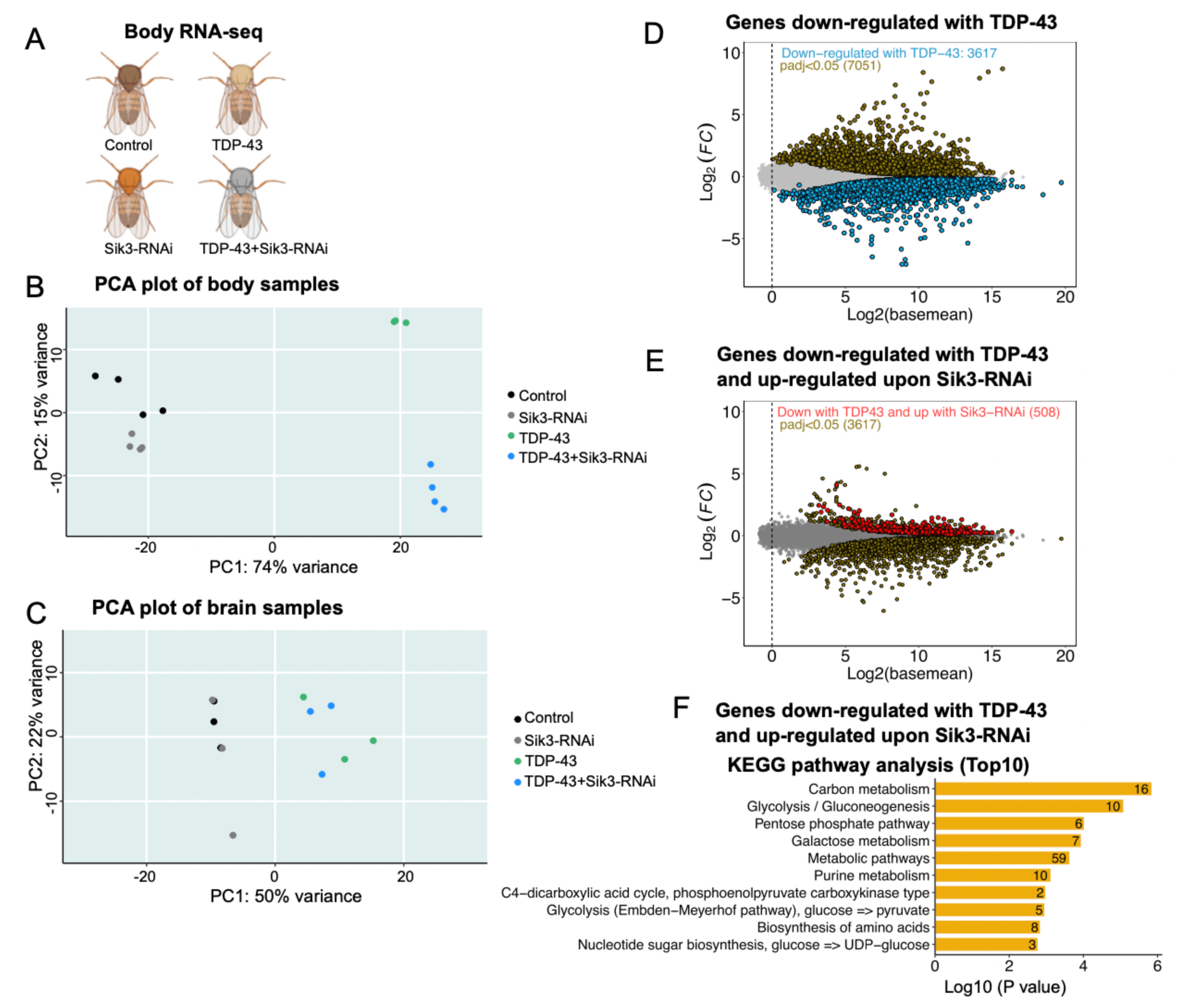
Enrichment of metabolic transcript recovery with Sik3 knockdown in bodies of TDP-43 flies. (**A**) RNA-seq experimental design. (**B**) Principal component analysis (PCA) of body samples. (**C**) PCA of brain samples. (**D**) MA plot of genes altered in TDP-43-expressing bodies. Gold marks all differentially expressed genes Padj < 0.05 (7051 genes), blue overlay highlights significantly down-regulated transcripts (3617). (**E**) MA plot of genes differentially expressed in bodies of animals expressing TDP-43 and Sik3-RNAi. Gold indicates all differentially expressed genes (3617) Padj < 0.05; red indicates genes that were significantly down-regulated with TDP-43 but now up-regulated with Sik3-RNAi. (**F**) KEGG pathway analysis of genes significantly down-regulated with TDP-43 and up-regulated with Sik3-RNAi identifies metabolic pathways, including glycolysis and pentose phosphate pathway. Genotypes: daGS; UAS-YFP x Luc-RNAi, daGS; UAS-TDP-43 x Luc-RNAi, daGS; UAS-YFP x Sik3-RNAi or UAS-TDP-43 x Sik3-RNAi, 6d on RU486.

Consistent with these PCA patterns, Sik3 RNAi selectively modified only a subset of TDP-43–dependent transcriptional changes in bodies. Of 3,617 genes downregulated by TDP-43, 508 were increased in the TDP-43 + Sik3 RNAi condition (Fig. 6D,E; Fig S7A,B). Pathway analysis of this subset revealed enrichment for metabolic processes, including glycolysis/gluconeogenesis, carbon metabolism, and the pentose phosphate pathway (PPP) (Fig. 6F). Importantly, many metabolic pathways remained enriched among the larger set of TDP-43–downregulated genes that were not corrected by Sik3 (Fig. S7C), indicating that Sik3 does not broadly normalize metabolic transcription but instead modulates specific metabolic components.

Comparison with Sik3 RNAi alone further indicated that the effects of Sik3 reflect partial overlap between its intrinsic transcriptional program and TDP-43–induced changes: approximately half of the genes normalized by Sik3 knockdown in the TDP-43 background were also upregulated by Sik3 RNAi alone (Fig. S7B). Together, these findings indicate that Sik3 reshapes, rather than reverses, the TDP-43 peripheral transcriptional landscape, selectively engaging metabolic pathways that are consistent with its ability to rescue sleep without globally restoring energy stores.

### Spatial metabolomics reveals selective restoration of peripheral PPP-associated metabolites by Sik3 knockdown

To determine whether the transcriptional signatures above corresponded to spatially resolved metabolic changes *in vivo*, we performed DESI-MSI–based spatial metabolomics^41,42^ on fly bodies and brains across genotypes (Fig. 7A). Similar to the RNAseq results, principal component analysis revealed strong genotype-driven separation in peripheral tissues. In bodies, TDP-43 samples separated from controls along PC1, and TDP-43 + Sik3 RNAi formed a distinct cluster rather than reverting fully to control (Fig. 7B). In contrast, brain profiles showed less pronounced separation across genotypes (Fig. S8A). Principal component analysis of individual metabolites (Fig. 7C) in the periphery showed TDP-43 expression leads to high levels of 6-Phosphogluconate and NADPH, among others (Fig. 7C). These changes implicate altered PPP activity in these flies, consistent with the observed transcriptional signatures. To further assess PPP pathways, we compared key PPP intermediates, including sedoheptulose-7-phosphate, ribose-5-phosphate, and xylulose-5-phosphate, and found these were altered in TDP-43 bodies and directionally shifted toward control levels in TDP-43 + Sik3 RNAi animals (Fig. 7D-F). Importantly, Sik3 knockdown alone had minimal effects on these metabolites (Fig. 7D-F), reinforcing that these changes are specific to the TDP-43 background. Analogous analyses in the brain revealed a different pattern (Fig. S8): while some metabolites were altered by TDP-43 expression, PPP-associated intermediates were not consistently affected, nor were they selectively normalized by Sik3 RNAi. Thus, the metabolic rescue signature associated with Sik3 appears to be predominantly peripheral, paralleling the RNA-seq profiles. The divergence between brain and peripheral metabolic signatures suggests that systemic metabolic state, rather than brain-intrinsic metabolic changes alone, is a dominant driver of sleep disruption in this model.

**Figure 7.**
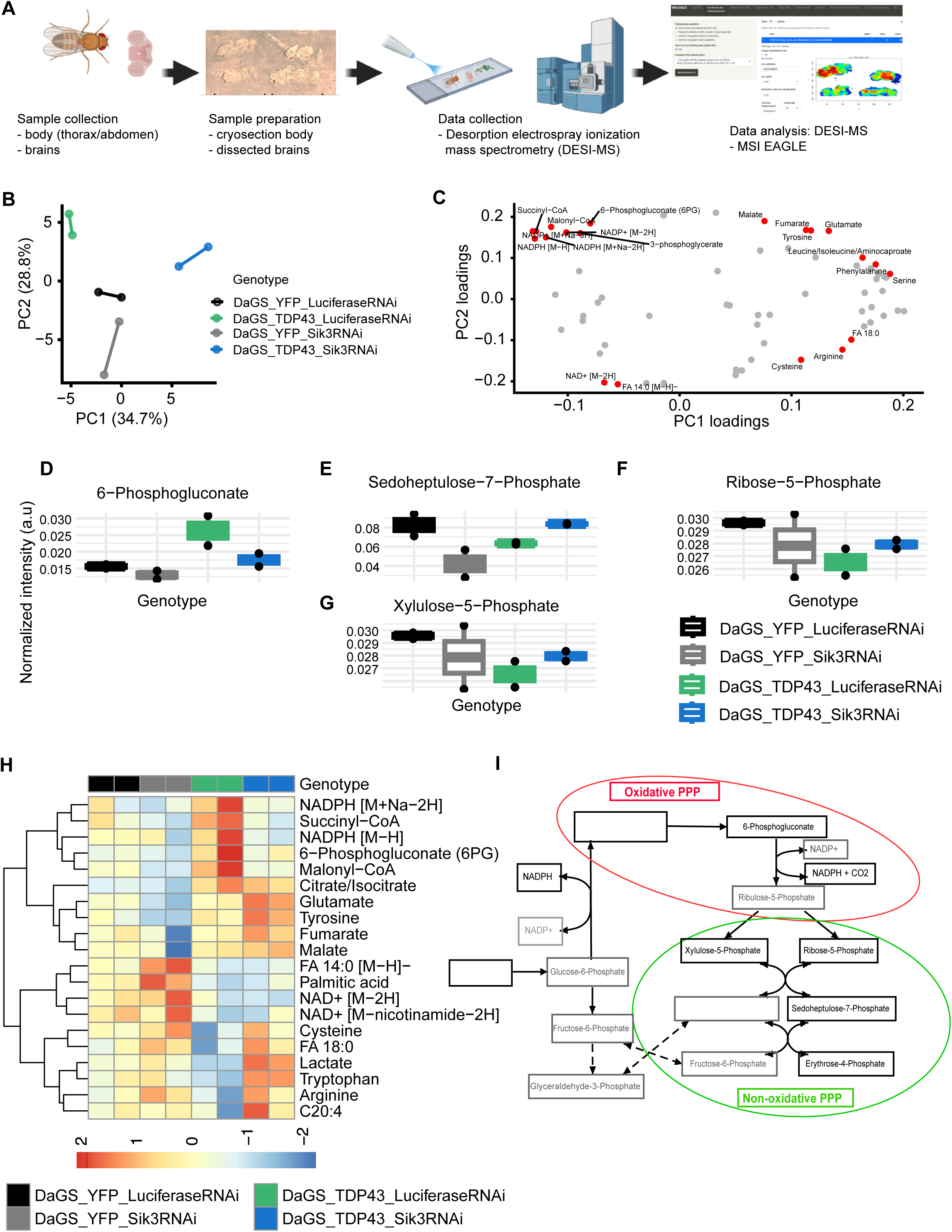
DESI–mass spectrometric imaging reveals alterations in the pentose phosphate pathway and associated metabolites in bodies of TDP-43 flies. (**A**) Schematic overview of data acquisition using desorption electrospray ionization mass spectrometric imaging (DESI-MSI). (**B**) Principal component analysis (PCA) scores plot based on the global mass spectral profiles of whole fly bodies, showing separation along the first two principal components. (**C**) PCA loadings plot highlighting the top 20 metabolites contributing to group separation (marked in red). (**D–F)** Boxplots depicting relative abundance of key pentose phosphate pathway (PPP) intermediates across four genotypes. (**G**) Heatmap of the top 20 metabolites identified in the PCA loadings plot (panel C). (**H**) Schematic representation of the pentose phosphate pathway; metabolites not detected by the DESI-MSI method are indicated in grey.

Interestingly, despite normalization of PPP-associated metabolites, transcriptomic data showed that expression of the PPP rate-limiting enzyme Zw (G6PD) remained reduced in bodies of TDP-43 flies expressing Sik3 RNAi, indicating that Sik3 does not rescue PPP function by restoring G6PD transcription (Fig. S9). By contrast, the carbohydrate-responsive transcription factor Mondo^43^ — upregulated in TDP-43 flies — was restored toward control levels with co-expression of Sik3 RNAi (although remained unaffected with Sik3 knockdown alone) (Fig. S9). These data are consistent with the recovery of a broader sugar-sensing and metabolic regulatory program, rather than the simple correction of a single enzymatic bottleneck.

Beyond canonical PPP intermediates, TDP-43 expression was also associated with high levels of succinyl, malonyl-CoA, and citrate, along with low levels of NAD+, free fatty acids (FA), and lactate (Fig. 7C,G). These observations suggest that elevated 6-phosphogluconate is not a result of overactive glycolysis; instead, the PPP oxidative arm appears activated at a functional level, consistent with redox or substrate-driven regulation. Low levels of pentose phosphate intermediates suggest oxidative PPP activation is primarily due to NADPH generation rather than nucleotide production. Simultaneous elevation of citrate and malonyl-CoA with reduced NAD+, lactate, and free fatty acids suggests bottlenecks in the TCA cycle and lipogenesis. Taken together, this pattern is consistent with a metabolic state suggestive of altered redox balance and constrained oxidative metabolism. Moreover, while the PPP metabolites were restored by Sik3 knockdown, TCA cycle metabolites remained elevated and NAD+ remained decreased (Fig. 7G). Thus, our findings suggest that Sik3 does not broadly restore peripheral metabolism to rescue sleep, but rather selectively modifies a PPP/redox-linked metabolic axis that has been disrupted by TDP-43 (Fig. 7H).

## Discussion

Sleep disruption in neurodegenerative disease is widely attributed to dysfunction within the brain^1,44^, yet the contribution of broader, non-neural physiological processes to disease-associated sleep phenotypes remains poorly defined. Here, we demonstrate that TDP-43 expression induces a chronic starvation-like state characterized by depletion of peripheral carbohydrate stores despite unchanged feeding, and that this metabolic state is sufficient to drive profound sleep loss. Restoring sleep alone fails to normalize peripheral metabolism, whereas improving metabolic state though dietary glucose availability rescues sleep, arguing that sleep disruption in this context reflects a downstream consequence of systemic metabolic imbalance. These findings support the idea that sleep phenotypes in neurodegenerative disease do not arise purely from dysfunction in the nervous system and that peripheral metabolic state plays a causal role.

A growing body of work across species indicates that TDP-43 disrupts cellular metabolism^10,45,46^, although the emphasis has largely been on neuronal or CNS-intrinsic effects. In *Drosophila* and rodent models, TDP-43 has been linked to altered glucose utilization, mitochondrial dysfunction, oxidative stress, and impaired energy balance; in humans with ALS and FTD, systemic metabolic abnormalities—including hypermetabolism, weight loss, and altered lipid profiles—predict disease trajectory^9,10,45,46^. Emerging studies have implicated redox and pentose phosphate pathway–related changes in models of ALS and other neurodegenerative disorders, consistent with a role for NADPH-dependent buffering and biosynthetic pathways in disease vulnerability^10,47,48^. However, most prior work has not addressed whether such metabolic alterations are functionally linked to behavioral perturbations such as sleep, or whether peripheral metabolic dysfunction is sufficient to drive sleep disruption independent of overt neuronal injury. Our results demonstrate that peripheral metabolic state, rather than brain impairment alone, can govern sleep behavior in TDP-43 proteinopathy.

SIK3 has emerged as a conserved regulator of sleep across species, yet its functional role appears context-dependent^19,40^. In mammals, gain-of-function mutations in SIK3 increase sleep, while loss-of-function reduces sleep, consistent with a role in baseline sleep homeostasis^40^. In contrast, studies in *Drosophila*^49^ and *C. elegans*^19^ suggest a more nuanced function, with SIK signaling linked to metabolic state and stress-responsive sleep-like behaviors rather than constitutive control of sleep amount. In our study, Sik3 knockdown alone does not produce a strong baseline sleep phenotype, yet robustly suppresses sleep loss in the context of TDP-43 expression. The nature of this metabolic stress signal may be informative: prior work in *Drosophila* has shown that SIK3 regulates the NADPH/NADP⁺ redox balance in the context of sugar tolerance^38^, and our metabolomic data show that TDP-43 expression elevates NADPH and 6-phosphogluconate while Sik3 knockdown normalizes these PPP-associated metabolites. This raises the possibility that SIK3 senses or responds to a redox or PPP-linked metabolic signal specifically, rather than to bulk energy depletion, and that it is this axis — rather than glucose or glycogen availability — that is transduced into sleep regulatory output.

The convergence of SIK3, a canonical sleep gene, with TDP-43, a defining proteinopathy in neurodegenerative disease, brings together sleep regulation and disease biology at a genetic level. Although prior work has largely focused on neuronal roles for SIK3^50^, its ability to suppress both sleep loss and starvation sensitivity in TDP-43 flies highlights a role in coordinating systemic metabolic programs relevant to sleep.

Consistent with this idea, transcriptomic and spatial metabolomic analyses show that SIK3 suppression does not restore bulk energy stores like glucose or glycogen, but instead selectively reshapes peripheral metabolic pathways associated with PPP-linked and redox-related metabolism. The dissociation between persistent suppression of central PPP enzyme transcripts and normalization of PPP-associated metabolites suggests that metabolic re-routing, rather than restoration of energy stores *per se*, underlies sleep rescue.

How is the peripheral metabolic state communicated to sleep-regulatory processes? One possibility is that chronic peripheral energy stress alters circulating metabolic signals or hormonal cues that directly influence sleep circuit neuronal excitability and arousal systems. Alternatively, peripheral redox imbalance may impose a systemic stress state that secondarily impairs brain metabolic resilience, which has already been implicated in sleep control. These models are not mutually exclusive, and it is possible that peripheral and central metabolic changes are not tightly coupled at the molecular level. Our findings support the idea that sleep disruption can arise from distinct metabolic perturbations — neuronal or peripheral — that converge on shared behavioral outputs. Such peripheral-driven sleep disruption may then feed forward to negatively impact brain health, consistent with evidence that chronic sleep loss exacerbates proteostatic stress, impairs metabolic homeostasis, and accelerates neurodegenerative processes^31,51,52^.

These findings raise additional questions to be addressed, including identifying signals that couple peripheral metabolic state to sleep-regulatory mechanisms and determining how peripheral metabolic interventions influence long-term neurodegenerative trajectories. At the same time, these results have intriguing implications for understanding altered sleep in neurodegenerative diseases. Sleep disturbances in ALS and FTD may, in part, reflect treatable systemic metabolic dysfunction rather than irreversible neuronal damage. Targeting peripheral metabolic pathways could therefore improve sleep and potentially mitigate downstream disease consequences, even without directly modulating brain pathology.

## Materials and Methods

### Fly strains

*Daughterless*-GS; UAS-*TDP43^52S^* flies were obtained from laboratory stocks. UAS-RNAi lines were sourced from the Bloomington Drosophila Stock Center. Flies were reared in bottles on standard molasses media (Lab Express) at 25°C with a 12:12 light-dark cycle. Animals were raised on this medium until adulthood, at which point they were placed on a 5% sucrose - 2% agar media supplemented with appropriate chemicals according to the different experiments. sik3-RNAi #1 is 56941, #2 is 28366, #3 is 57302, we used 57302

### RU486 food preparation

RU486 (Thermo Fisher Scientific) was prepared as a 50 mM stock solution in 100% ethanol. Molasses media or sucrose-agar media was melted, and appropriate volume of RU486 or ethanol was added and mixed once cooled to make a final concentration of 0.5 mM. This molten mixture was then poured into empty polystyrene vials (Genesee Scientific) or used to fill glass DAM tubes (Trikinetics 6x15).

### Gaboxadol

As in RU486 protocol, gaboxadol was added to standard molasses fly media or sucrose agar media at a final concentration of 0.1 mg/ml.

### High-glucose diet (10x Sugar)

High-glucose media of 1.6% agar, 80% glucose, 8% inactive yeast, and 1.6% acid mixture (phosphoric acid + propionic acid) was prepared in distilled water and poured into empty polystyrene vials. For DAM tubes, media containing 50% glucose and 2% agar was used.

### Café assay

10 male flies were added to vials containing a 2% agar water solution. Vials were sealed with a CAFÉ stopper, and the flies were starved overnight, with an empty capillary tube placed to allow habituation to the feeding apparatus. The next morning, the empty capillary tube was replaced with one filled with a 2% sucrose-water solution supplemented with blue dye. Mineral oil was added at the end of the capillary to prevent evaporation. The vials were placed horizontally in an incubator at 25 °C with a 12:12 h light: dark cycle. The initial meniscus position of the food solution was marked at the start of the assay and food intake was recorded 6 hours later as the change in meniscus position. Flies that died after the overnight fast were excluded, and the amount of food intake was divided by the number of living flies in the vials.

### Sleep analysis

Male flies were collected at 1-3 days post-eclosion under brief CO2 anesthesia and housed in groups. After 2 days, individual flies were loaded into glass DAM tubes containing the appropriate diet and pharmacological supplements. For all experiments, data collection began at ZT0 following CO2 anesthesia, and the first 24 hours of data were discarded. Locomotor activity was monitored using the DAM system (Trikinetics) and recorded in 1-minute bin. Sleep was operationally defined as 5 minutes of consolidated inactivity. Data were processed with a custom R script using Rethomics package.

### Glucose & Glycogen Assay

Whole-fly body glucose and glycogen levels were quantified using the Glucose (GO) assay kit (Sigma-Aldrich) following a published protocol^53^. Male flies 1-3 days post-eclosion were collected and maintained for 6 days in vials with their assigned diet conditions. Afterwards, they were flash-frozen on dry ice and vortexed at high speed to separate heads from bodies. Bodies were collected on ice and homogenized in cold PBS (5 bodies in 125 µL of PBS) using a motorized pestle. An aliquot of the homogenate was removed at this time to measure protein content with a Pierce BCA Protein Assay (Thermo Fisher Scientific) and stored at -80°C for later analysis. The remaining supernatant was heat-treated for 10 minutes at 70°C, then centrifuged and transferred to a new microfuge tube. Supernatants were diluted 1:5 in PBS and stored at -80°C for later analysis. Glucose standards were prepared according to the protocol and plated alongside samples in a 96-well hard-shell PCR plate with clear wells (Bio-Rad). Reagents were added as per kit instructions. Absorbance was measured at 540 nm using a microplate reader. Glucose concentration was determined using a standard curve and normalized to the total protein content in each sample.

For glycogen concentration, a 1mg/ml glycogen stock solution was prepared and serially diluted to generate glycogen standards in parallel with glucose standards. Each sample was plated in duplicate: one to determine free glucose concentration, the other to determine glycogen content after enzymatic digestion with amyloglucosidase (Sigma A1602). To determine glycogen concentration, the absorbance measured in the free glucose samples was subtracted from the absorbance measured in the amyloglucosidase-digested samples. The glycogen content was then determined based on the glycogen standard curve and normalized to protein.

### Starvation protocol

For starvation experiments, male flies were collected and maintained the same as for sleep analysis. Following a day of recording baseline sleep, they were transferred into DAM tubes containing only 2% agar and RU486 or EtOH, to provide hydration without nutrients. Locomotor activity was monitored using the DAM system (Trikinetics). Data were processed using a custom R script to extract time-to-death for each individual fly. Survival data were then plotted as Kaplan-Meier survival curves and analyzed on GraphPad Prism.

### Whole-Animal Respirometry

Whole-animal indirect calorimetry was performed using a flow-through respirometry platform based on the MAVEn system to measure carbon dioxide production (VCO₂), with oxygen consumption (VO₂) measured downstream using an Oxzilla differential oxygen analyzer, as described previously^54^. Male flies were collected 1-3 days post-eclosion and maintained for 6 days at 25 °C on standard molasses-based food containing either RU486 or ethanol (EtOH) control. On day 6, flies were briefly anesthetized on ice and loaded into glass respirometry chambers, with 25 flies per condition. Each chamber was supplied with 2 mL of food consisting of 5% sucrose and 2% agar supplemented with RU486 or EtOH as appropriate. Control chambers containing identical food, but no flies were included to account for background signal. During the recording period, air was passed continuously through the chambers and directed sequentially through the gas-analysis system for measurement of CO₂ production and O₂ consumption. The initial hours after chamber loading were excluded from analysis to minimize handling-related disturbance and allow metabolic stabilization. Scrubbing columns were replaced daily throughout the experiment to maintain signal quality and analyzer performance. Respiratory quotient (RQ) was calculated as the ratio of CO₂ produced to O₂ consumed and used as a measure of substrate utilization.

### RNAi-based genetic screen

Virgins collected from *Daughterless*-GS>UAS-*TDP43^52S^*stock were crossed to males of RNAi lines. An mCherry RNAi line was used as a negative control. Male flies (12-16 per RNAi line) were collected and loaded onto RU486 DAM tubes (or ethanol vehicle, where indicated). Sleep experiments were performed as described above.

### Western immunoblotting

Body samples were homogenized in LDS buffer composed of 325 µL 4× LDS buffer (Invitrogen, NP0007), 185.9 µL protease inhibitor stock solution (Roche, 11836170001), 13 µL 1 mM phenylmethylsulfonyl fluoride (Sigma-Aldrich, P7626), and 32.5 µL β-mercaptoethanol (Sigma-Aldrich, M6250). 7.5 μl of sample buffer is added per body. Samples are boiled at 95°C for 5 min and centrifuged at 1500 rpm for 3 min at room temperature. Samples were loaded onto 15- well 1.0- mm 4 to 12% bis-tris NuPAGE gels (Thermo Fisher Scientific, WG1401) with prestained protein ladder (Thermo Fisher Scientific, 22619). One head was loaded per lane. Gel electrophoresis was performed using Xcell Surelock Mini-Cell Electrophoresis System at 120 V. Proteins were transferred overnight onto a nitrocellulose membrane (Bio-Rad, 1620115), using a Bio-Rad mini transblot cell at 90 A for 17 hours. Membranes were stained in Ponceau S (Sigma-Aldrich, P7170-1L), washed with deionized water, and imaged using Amersham Imager 600. Ponceau S staining was removed by washing off 3 times for 5 min in Tris-buffered saline with 0.1% Tween 20 (TBST). Membrane was blocked in 5% nonfat dry milk (LabScientific, M08410) in TBST for 1 hour and incubated with primary antibodies with blocking buffer overnight at 4°C. After washing 3 times for 5 min in TBST, membrane was incubated with horseradish peroxidase-conjugated secondary antibodies for 2 hours at room temperature. Membrane was washed 3 times for 10 min in TBST, and signal was detected using ECL prime (Cytivia, RPN2232) and imaged with an Amersham Imager 600. Primary antibodies used were anti-tubulin (1:12,000; Developmental Studies Hybridoma Bank, #AA4.3) and anti–TDP-43 (1:10000; ProteinTech, #10782-2-AP). Secondary antibodies were goat anti-mouse (1:10000; Jackson lmmunoResearch, #115-035-146) and goat anti-rabbit (1:10000: Jackson lmmunoResearch, #111-035-144).

### RNA sample preparation

Fly brains or bodies were dissected after 6 days on RU486 food. n = 4 biological replicates, 10 to 12 brains per sample, 5 bodies (minus head) per sample. Tissue was homogenized in 200 μl of TRIzol (Thermo Fisher Scientific, 15596026) in ribonuclease-free 1.5-ml microfuge tubes (Thermo Fisher Scientific, AM12400). A total of 800 μl of TRIzol (Thermo Fisher Scientific, 15596026) was added to the tube, and 200 μl of chloroform (Thermo Fisher Scientific, AC423555000) was vigorously shaken for 20 s at room temperature. Samples were left for 5 min at room temperature to form upper aqueous phase and centrifuges at 4°C for 15 min at 12,000g. The upper aqueous phase was transferred to a fresh ribonuclease-free tube. RNA samples were then processed using the Zymo RNA clean & concentrator -5 kit (Zymo, R1013), using their RNA clean-up from aqueous phase after TRIzol/chloroform extraction protocol plus on column deoxyribonuclease I treatment.

### RNA-seq analysis

RNA-seq libraries were generated using the TruSeq stranded mRNA Library Preparation kit and sequenced by Admera Health on an Illumina NovaSeq S4 platform, yielding approximately 40 million paired-end reads per sample (2 x 150 bp). For each genotype four biological replicates were prepared. Raw paired-end fastqs files were processed with TrimGalore (v.0.6.7) (https://github.com/FelixKrueger/TrimGalore) with default settings to remove Illumina adapters. Trimmed pair-end reads were aligned to *Drosophila melanogaster* reference genome (r6.48) using HISAT2 (v 2.2.1) with a list of known splice sites. HISAT2 index was built from *D. mel* reference genome (r6.48). Reads that failed to align were not included in downstream data processing. Samtools (v 1.15.1) was used to convert SAM files to BAM files, sort alignments by coordinates, and index coordinate-sorted BAM files. Genes with low read counts (sum of reads in all samples less than 10) were filtered and not included in downstream analysis. Differential expression analysis was performed using DESeq2 (v.1.34.0). Principal components analysis (PCA) plots were made using the plotPCA function in DESeq2, with variance stabilized counts as the input. MA plots were constructed from the adjusted P-values and baseMean values output from DESeq2. Normalized counts produced by DESeq2 were used to show expression levels^55–58^.

### GO and pathway analysis

GO analysis and pathway enrichment analysis for differentially expressed genes were performed using FlyMine (v.53) and PANGEA (v.2). The test correction was set to Holm--Bonferroni with a maximum P-value of 0.05. A list of all genes with detectable expression was used as background for both GO and pathway analysis. Redundant GO terms were removed using Revigo (v 1.8.1).

### Desorption Electrospray Mass Spectrometry Imaging (DESI-MSI)

Tissue sections were suspended in 3.5% carboxymethylcellulose (CMC) media (C4888, Sigma Aldrich) and sectioned at a thickness of 10 μm using a Leica CM1950 cryostat. The sections were thaw-mounted onto superfrost glass slides and stored at -80°C until analysis. Mass spectrometry imaging (MSI) data were acquired using a Xevo G2-XS QTOF mass spectrometer equipped with a DESI-XS source (Waters Corporation, Milford, MA, USA). Data were acquired in negative ion mode between m/z 50-1200. The spray solvent consisted of 98% methanol with 50 pg/mL leucine enkephalin at a flow rate of 1 μL/min. The spray voltage was set to -0.6 kV, nebulizing N_2_ gas flow maintained at 600 L/H. For “small molecule” acquisitions, the collision cell RF offset was set to 150 V and heated transfer line temperature 50°C. The sampling area was defined using High Definition Imaging (HDI) v1.7 with pixel size 20 x 20 μm, and raster rate 80 μm/s.

### Mass Spectrometry Imaging Data Analysis

The .RAW continuum MSI data files were first lockmass corrected (leucine-enkephalin reference: m/z 554.2615) and centroided using the All File Accurate Mass Measure routine in MassLynx v4.2, converted to mzML format with msConvert^59^, and finally converted to imzML format using imzMLConverter v2.1.1 for analysis^60^.

Processing, statistical analysis, and visualization of MSI data was performed with MSI.EAGLE v0.1^61^. In MSI.EAGLE, spatial segmentation was carried out using Spatial Shrunken Centroids (SSC) analysis to distinguish tissue from background ionization. Image masks derived from this segmentation were applied to isolate signals originating exclusively from tissue regions. Tissue-specific peak-picking was then performed using targeted metabolite lists or the "diff" method to generate untargeted metabolite feature lists.

## Figure Legends

**Supplementary Figure 1.**
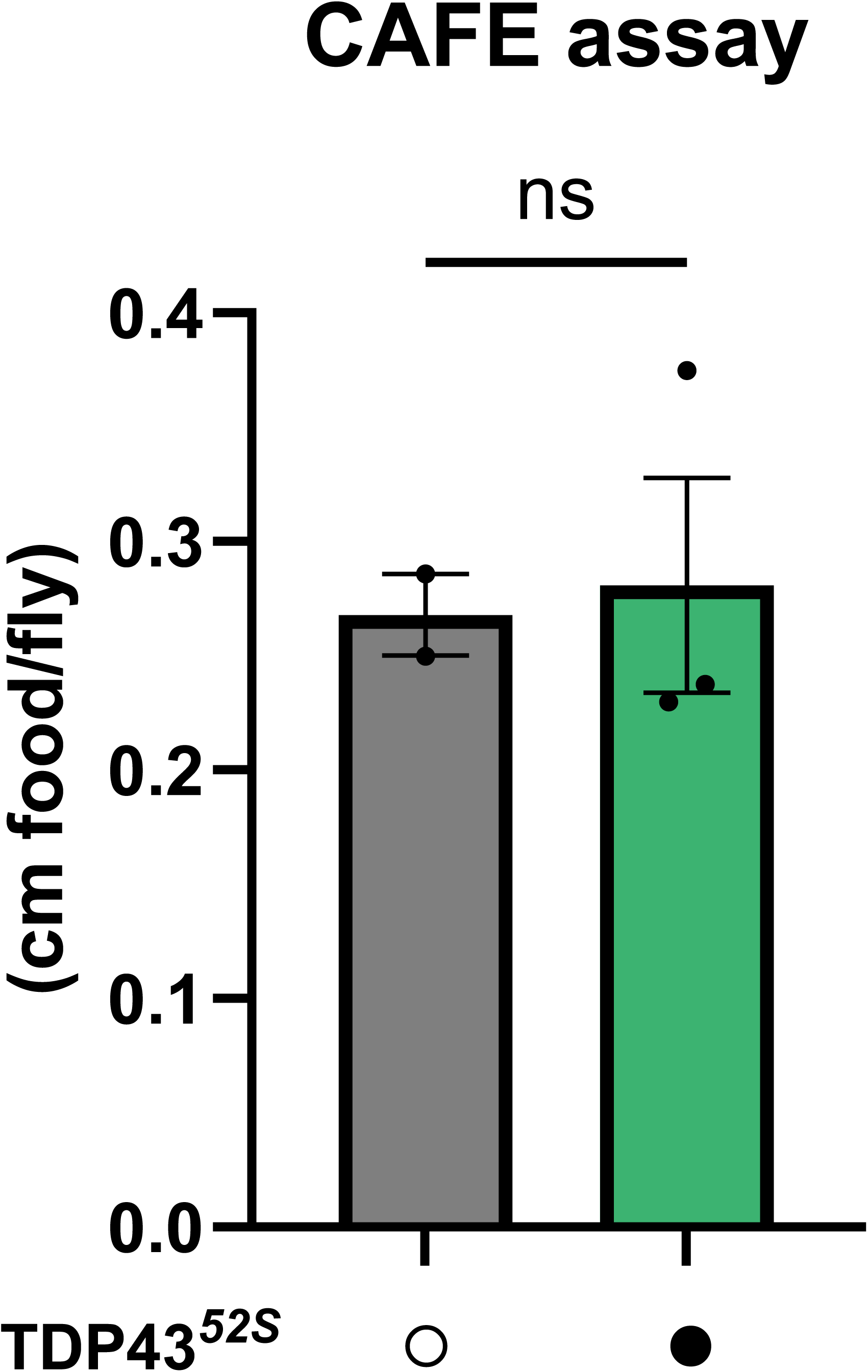
TDP-43 flies do not show major changes in food consumption. Food intake is the same between *TDP-43* flies (n=3 groups of 10 flies) and control flies (n=2 groups of 10 flies), Mann Whitney test.

**Supplementary Figure 2.**
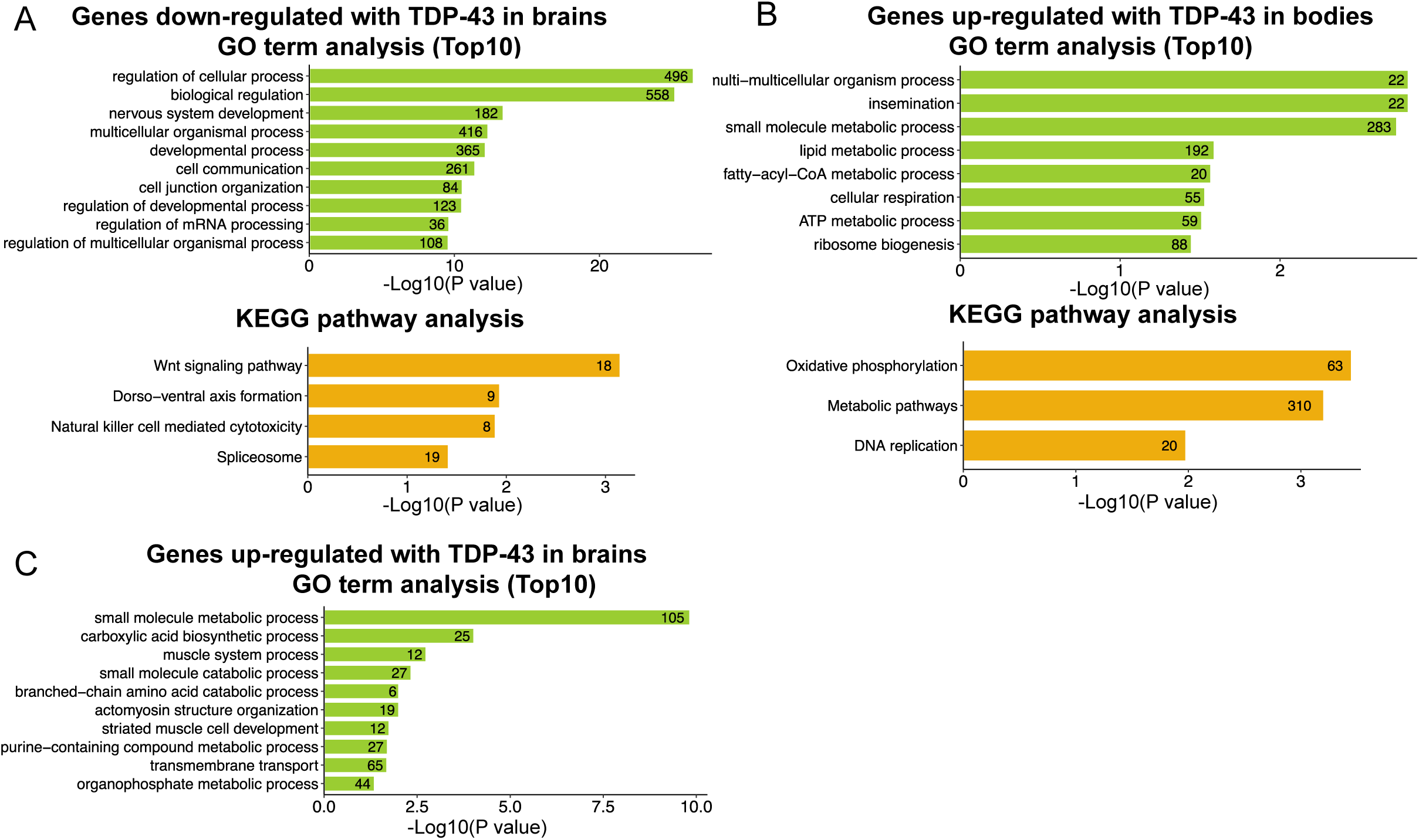
GO term and KEGG pathway analysis of DEGs from brains and bodies of TDP-43 animals. (**A**) GO term and KEGG pathway analysis of genes down-regulated in brains of TDP-43 flies. (**B**) GO term and KEGG pathway analysis of genes up-regulated in bodies of TDP-43 flies. (**C**) GO term analysis of genes up-regulated in brains of TDP-43 flies. Body genotypes: daGS; UAS-YFP x RNAi control or daGS; UAS-TDP-43 x RNAi control, 6d on RU486. Brain genotypes: daGS; UAS-YFP x RNAi control, daGS; UAS-TDP-43 x RNAi control, daGS; UAS-YFP x Sik3-RNAi or UAS-TDP-43 x Sik3-RNAi.

**Supplementary Figure 3.**
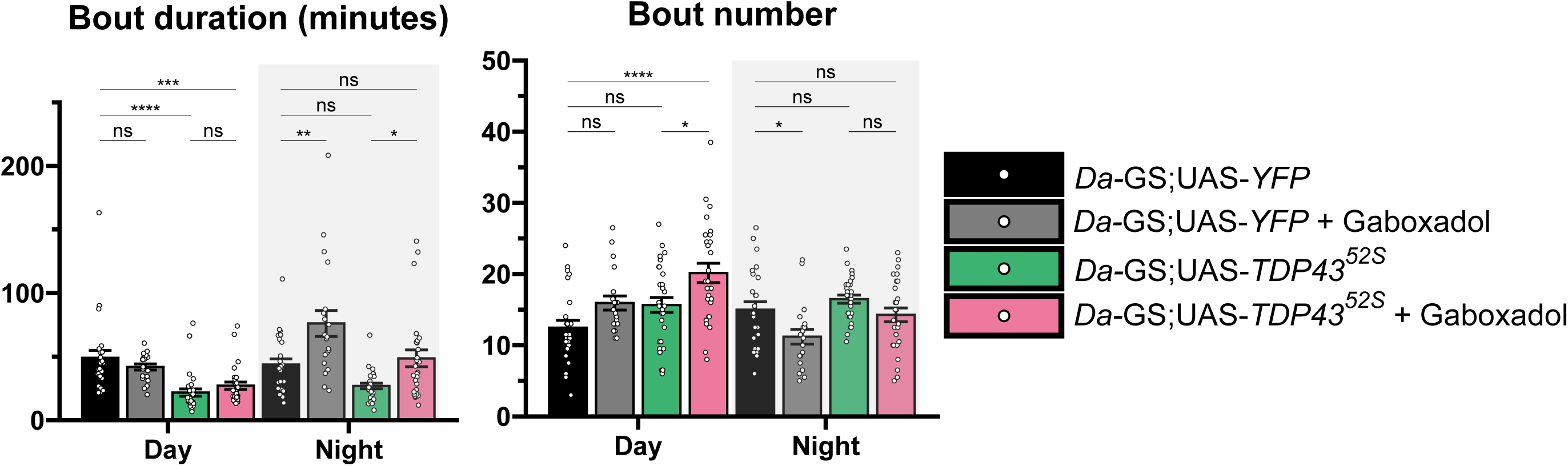
Sleep fragmentation measures related to. Figure 3. Sleep bout duration and number in TDP-43 flies treated with gaboxadol. From left to right: control flies (n=29 flies), control flies + gaboxadol (n=20 flies), *TDP-43* flies (n=25 flies), *TDP-43* flies + gaboxadol (n=27 flies). \**p* ≤ 0.05, \*\**p* ≤ 0.01, *** *p* ≤ 0.001, and \*\*\*\**p* ≤ 0.0001 by one-way ANOVA with Šídák’s multiple comparisons test.

**Supplementary Figure 4.**
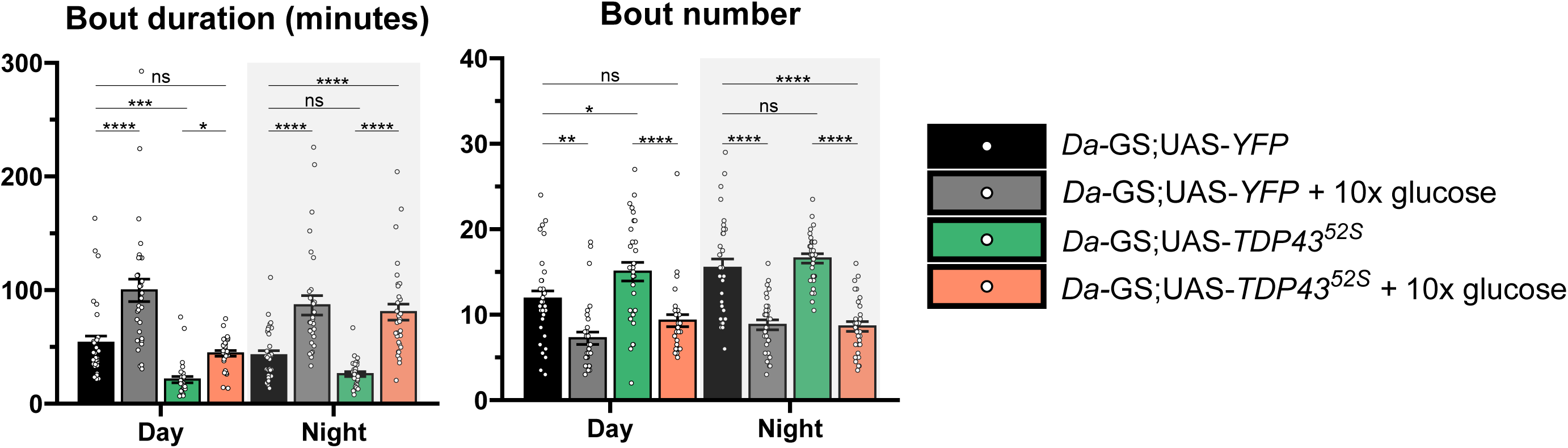
Sleep fragmentation measures related to. Figure 4. Sleep bout duration and number in TDP-43 flies treated with high glucose. From left to right: control flies (n=32 flies), control flies + 10x glucose (n=31 flies), *TDP-43* flies (n=31 flies), *TDP-43* flies + 10x glucose (n=32 flies). \**p* ≤ 0.05, \*\**p* ≤ 0.01, *** *p* ≤ 0.001, and \*\*\*\**p* ≤ 0.0001 by one-way ANOVA with Šídák’s multiple comparisons test.

**Supplementary Figure 5.**
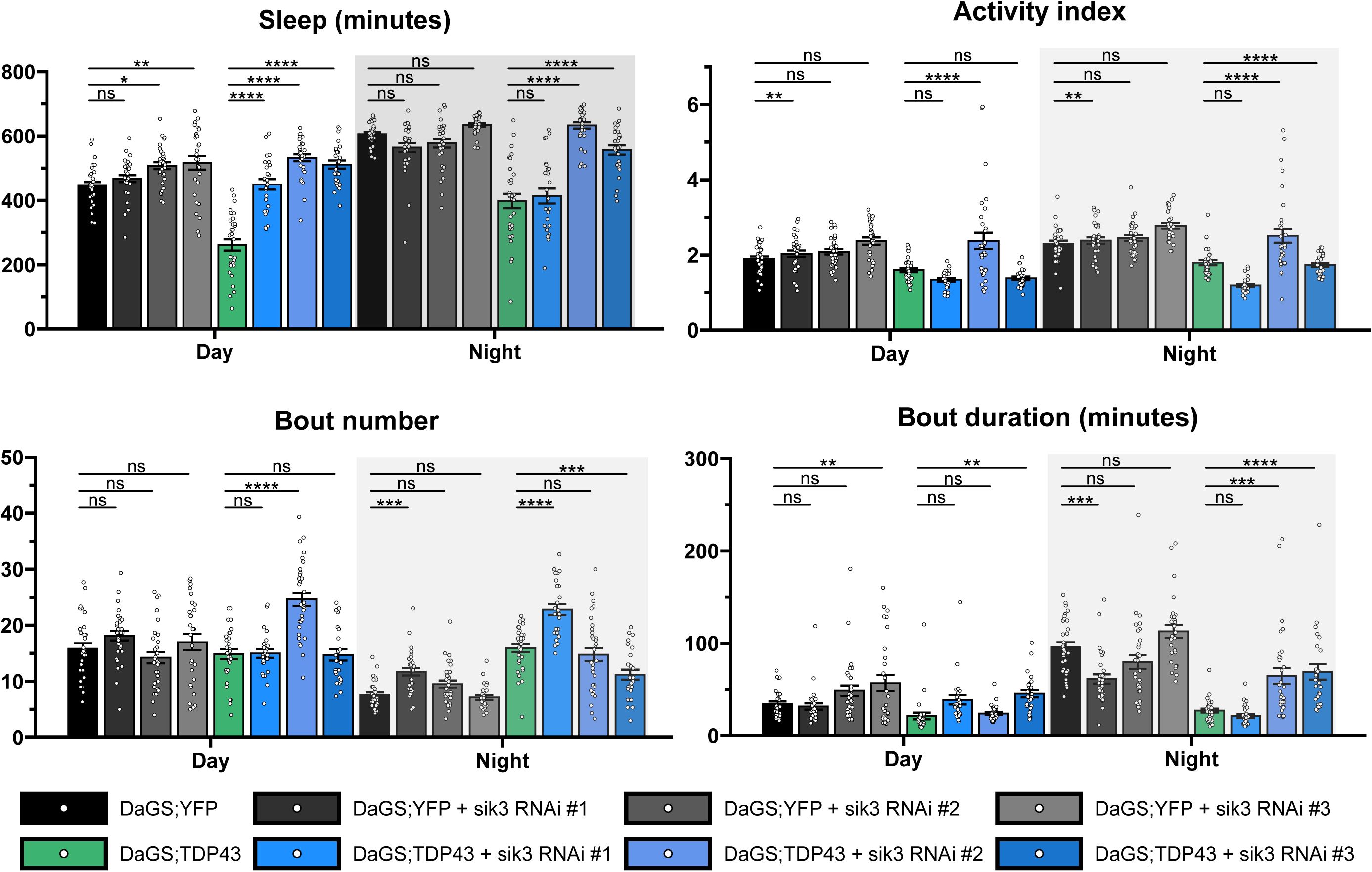
Sleep measures related to. Figure 5. Sleep duration, bout length, and number for three Sik3 RNAi lines. From left to right: control flies (n=31 flies), control flies + sik3 RNAi #1 (n=31 flies), control flies + sik3 RNAi #2 (n=32 flies), control flies + sik3 RNAi #3 (n=28 flies), *TDP-43* flies (n=31 flies), *TDP-43* flies + sik3 RNAi #1 (n=29 flies), *TDP-43* flies + sik3 RNAi #2 (n=32 flies), *TDP-43* flies + sik3 RNAi #3 (n=27 flies). \**p* ≤ 0.05, \*\**p* ≤ 0.01, *** *p* ≤ 0.001, and \*\*\*\**p* ≤ 0.0001 by one-way ANOVA with Šídák’s multiple comparisons test.

**Supplementary Figure 6.**
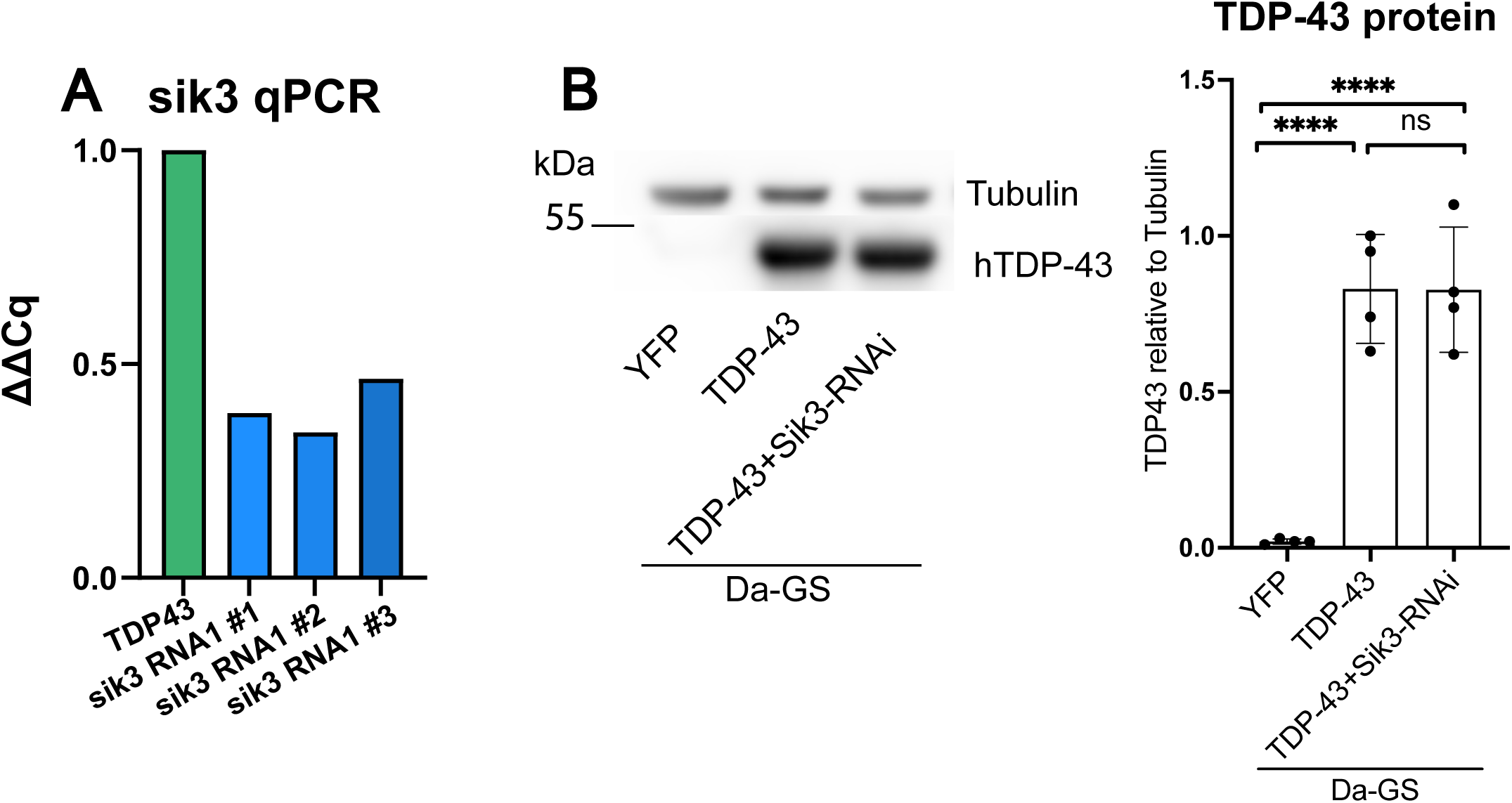
Sik3 validation studies. (**A**) qPCR confirming reduction of Sik3 in bodies of flies with RNAi expression under control of DaGS. (**B**) Western immunoblot and quantification of TDP-43 protein from fly bodies following 7 days on RU486.

**Supplementary Figure 7.**
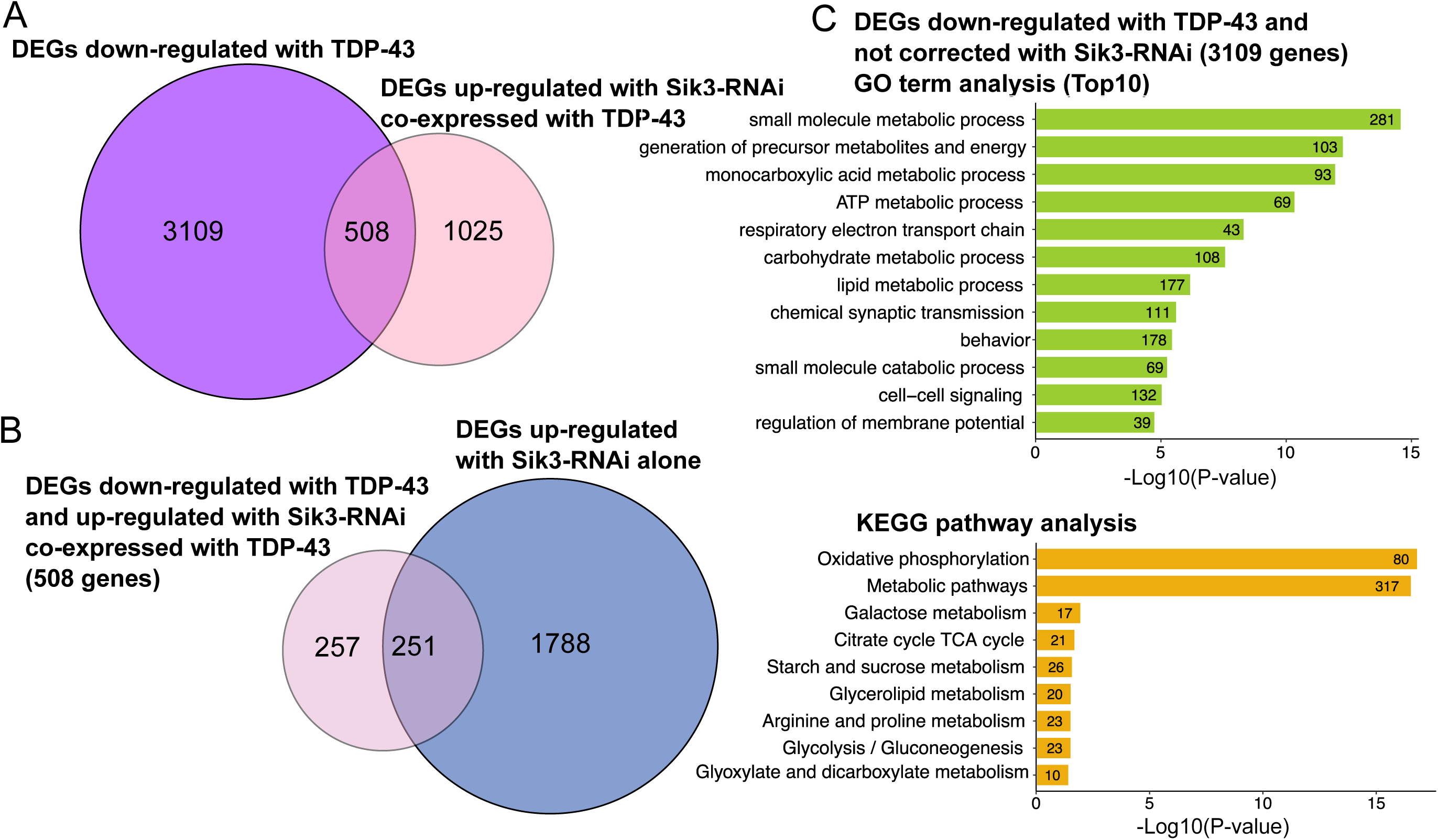
Sik3 knockdown partially restores TDP-43–dependent metabolic gene expression in fly bodies. (**A**) Venn diagram showing the overlap between transcripts down-regulated in bodies of TDP-43–expressing flies and transcripts up-regulated in TDP-43 flies upon Sik3 RNAi, identifying 508 genes whose expression is restored in the TDP-43 + Sik3 RNAi condition. (**B**) Venn diagram comparing the 508 restored transcripts with genes up-regulated by Sik3 RNAi alone, indicating that a substantial subset (251 genes) is also responsive to Sik3 knockdown in the absence of TDP-43.

**Supplementary Figure 8.**
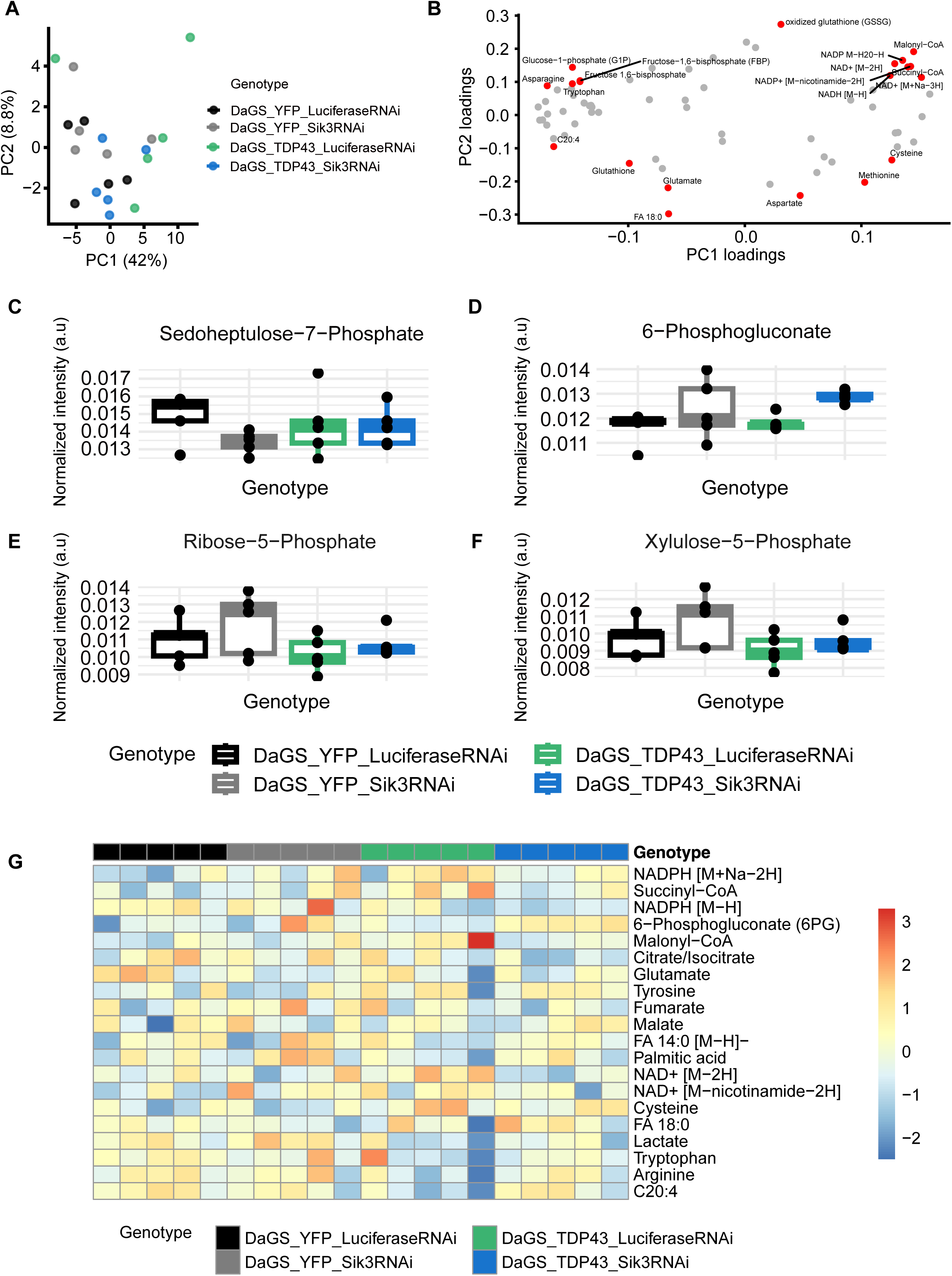
Brain metabolic profiles of flies analyzed by DESI–mass spectrometric imaging. (**A**) Principal component analysis (PCA) scores plot derived from global mass spectral profiles of fly brains, showing no clear separation among genotypes along the first two principal components. (**B**) PCA loadings plot highlighting the top 20 metabolites contributing to the variance captured by the first two principal components. (**C–E**) Boxplots showing the relative abundance of key pentose phosphate pathway (PPP) intermediates across four genotypes. (**F**) Heatmap of the top 20 metabolites identified in the PCA loadings plot of whole fly bodies (Figure 8, panel C), displayed here for comparison in brain tissue.

**Supplementary Figure 9.**
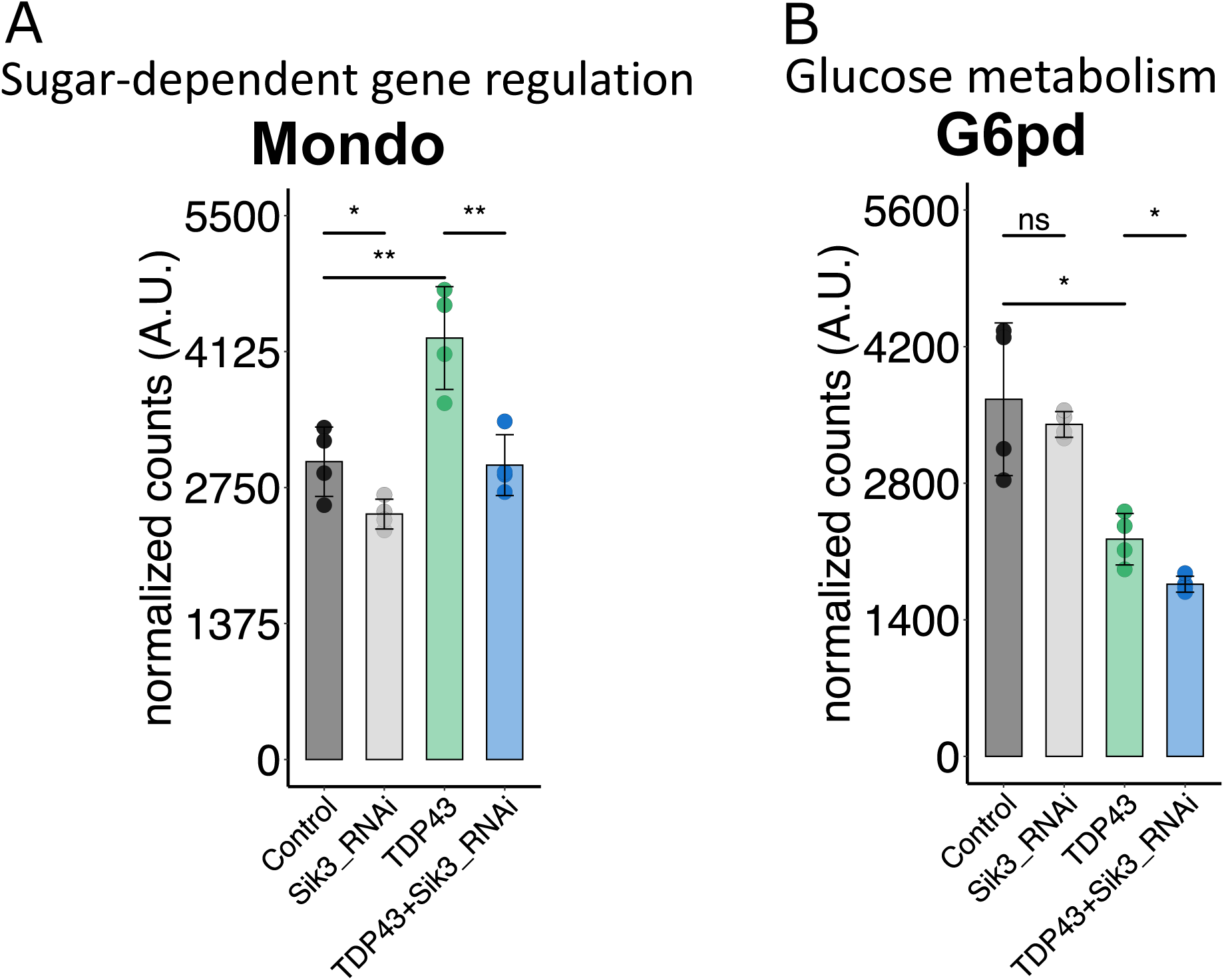
Expression of sugar-responsive and PPP-related genes in bodies of TDP-43 and Sik3 RNAi flies. Normalized RNA-seq counts for Mondo (left), a carbohydrate-responsive transcription factor, and G6pd (Zw) (right), the rate-limiting enzyme of the pentose phosphate pathway, across indicated genotypes. TDP-43 expression increases Mondo expression, which is partially restored toward control levels by co-expression of Sik3 RNAi. G6pd expression is reduced in TDP-43 flies and is not rescued by Sik3 knockdown. Each data point represents an independent biological replicate. Bars represent mean ± SEM. *p ≤ 0.05, **p ≤ 0.01, one-way ANOVA with Šídák’s multiple comparisons test.

## Acknowledgments

We thank Dr. Tania Reis, members of the Kayser Lab, Raizen Lab, Dr. David Raizen, and members of the Penn Chronobiology and Sleep Institute for helpful discussion and input.

## Author contributions

Conceptualization: MSK, NMB Investigation: All authors Writing – Original Draft: MSK Writing – Review and Editing: All authors Project Supervision and Funding: MSK, NMB

## Data and materials availability

All data needed to evaluate the conclusions in the paper are present in the paper and/or the Supplementary Materials.

